# Immune surveillance at plasmodesmata restricts the movement of a fungal effector

**DOI:** 10.64898/2026.05.19.726166

**Authors:** Emma K. Turley, Daniel S. Yu, Andrew Breakspear, Joanna Jennings, Mark Jave A. Bautista, Sally Jones, Barkha Ravi, Rafał Zdrzałek, Weibin Ma, Adam R. Bentham, Lauren S. Ryder, Nicholas J. Talbot, Mark J. Banfield, Christine Faulkner

## Abstract

To establish infection, phytopathogens deploy effectors to compromise host defences and facilitate invasive growth. As part of this, the battle for control of symplastic connectivity via plasmodesmata is a key determinant of infection outcomes, yet little is known about how fungal effectors directly exploit these channels, and in turn, how hosts defend them. Here, we have identified ChEC108 as a plasmodesmal-targeting, cell-to-cell mobile effector from the anthracnose fungus, *Colletotrichum higginsianum*. ChEC108 binds the plasmodesmal protein HEAVY METAL-ASSOCIATED (HMA) ISOPRENYLATED PLANT PROTEIN 6 (HIPP6) from Arabidopsis via a tetrahedral metal ion coordination site with either of its HMA domains. This binding event induces plasmodesmal closure, impairing the cell-to-cell mobility of ChEC108. Alongside the finding that loss of either *HIPP6* from the host or *ChEC108* from *C. higginsianum* favours infection, we conclude that HIPP6 detects the presence of an invading fungus and elicits defence.

## Introduction

Pathogens deploy diverse mechanical and chemical mechanisms to invade plant tissues and establish an infection, including the delivery of suites of effectors into host tissues. Pathogen effectors target a variety of host processes, often to enhance the infection process and gain control of host cells but can also be detected by immune receptors that activate defence upon effector perception. Plasmodesmata, the cytoplasmic connections between plant cells, offer a direct route for molecules, and pathogens themselves, to access uninfected cells and expand their invasion, making these channels prime targets for manipulation by effectors. Plants defend their plasmodesmata during pathogen invasion; extracellular perception of bacterial and fungal elicitors triggers localised signalling cascades and deposition of callose (1–3) to restrict plasmodesmal aperture (4) and isolate infected cells. As both the host and pathogen invader regulate these structures, plasmodesmata have emerged as a key site at which molecular plant-microbe interactions can influence infection and immune success.

The fungal anthracnose pathogen, *Colletotrichum higginsianum,* infects Brassicaceous plants including widely cultivated crops and the model plant, *Arabidopsis thaliana* (5, 6). As part of its hemibiotrophic lifecycle, the fungus penetrates host cells using a melanised appressorium and subsequently enters a biotrophic phase during which invasive hyphae proliferate within living host cells (6). Shortly after, fungal development undergoes a switch to necrotrophy, involving death and degradation of host tissues as thin filamentous hyphae radiate from initially penetrated cells. Collectively, the interactions of secreted effectors with extra-and intracellular host components are thought to be key determinants of *C. higginsianum* virulence on any given species. The genome of strain IMI 349063 was initially predicted to encode a repertoire of 365 candidate effectors (7) and analysis of expressed sequence tags by Kleemann et al. (8) indicated 198 Effector Candidate (*ChEC*) genes are transcribed during i*n planta* infection stages. Screening in *Nicotiana benthamiana* indicated *C. higginsianum* effectors can target multiple subcellular compartments in plants including the cytoplasm, nuclei, peroxisomes, and microtubules (9, 10), and while many can move between cells in plant tissues (9), plasmodesmal effectors have yet to be identified. Furthermore, very few *C. higginsianum* effectors have been functionally characterised and their specific host targets remain unknown.

In the last decade, it has become increasingly apparent that HEAVY METAL-ASSOCIATED (HMA) domain-containing proteins, including HMA PLANT PROTEINS (HPPs) and HMA ISOPRENYLATED PLANT PROTEINS (HIPPs) (11), are common host targets of phytopathogen effectors. For instance, *Magnaporthe* AVRs and ToxB-like (MAX) family effectors have been shown to target the HMA domains of HPPs or HIPPs from cereals infected by the blast pathogen, *Magnaporthe oryzae* (12–17). Additionally, a viral movement protein (18) and effectors from other fungal (19) and nematode (20) pathogens bound HMA proteins from their respective host plants. In turn, some plant species have evolved to exploit HMA domains for effector recognition, integrating them as baits into nucleotide-binding, leucine-rich repeat (NLR) (21, 22) and kinase fusion protein (KFP) (23, 24) receptor architectures. These discoveries suggest host HMA proteins have widespread roles in plant immunity that diverse pathogens have convergently evolved to exploit.

In this study, we report the interaction of *C. higginsianum* effector ChEC108 with the Arabidopsis host protein HIPP6, both of which locate at plasmodesmata. Binding of ChEC108 occurs via the HMA domains of HIPPs and depends on metal ion co-ordination. When HIPP6 binds ChEC108, plasmodesmal closure is induced, restricting the cell-to-cell movement of ChEC108 and impeding *C. higginsianum* infection. Therefore, our data suggests HIPP6 functions as an immune surveillance protein that recognises a fungal effector to activate immunity.

## Results

### The fungal effector ChEC108 targets plasmodesmata and moves cell-to-cell

In a screen for the host sub-cellular localisation of *C. higginsianum* effectors expressed during the biotrophic phase of infection (9), we identified ChEC108 (CH63R_09563) as a candidate effector that targets plasmodesmata. Excluding the predicted 21-amino acid N-terminal signal peptide (SP), an eGFP fusion to the C-terminus of ChEC108 localised to both nuclei and plasma membrane-associated puncta, the latter of which co-localised with aniline blue-stained plasmodesmal callose deposits when transiently expressed in *N. benthamiana* leaves (Fig. 1A). To confirm that ChEC108 can be translocated into host cells during infection, we assayed ChEC108 secretion from *Magnaporthe oryzae* and uptake into rice cells. At 26-30 hours post-inoculation, SP-ChEC108-eGFP signal was concentrated at a single punctum associated with each invasive hypha (*SI Appendix,* Fig. S1A), corresponding to the biotrophic interfacial complex (BIC) - a membrane-rich structure into which cytoplasmic *M. oryzae* effectors are secreted before translocation (25). With the SP removed, ChEC108-eGFP remained diffusely localised within *M. oryzae* hyphae and appressoria (*SI Appendix,* Fig. S1B). When *SP-ChEC108-eGFP-NLS* (including a non-native nuclear localisation signal) was expressed in *M. oryzae*, fluorescent signal was observed at the BIC and within host cell nuclei (*SI Appendix,* Fig. S1C), demonstrating that ChEC108 can be translocated from a fungus to plant cells.

**Fig. 1.**
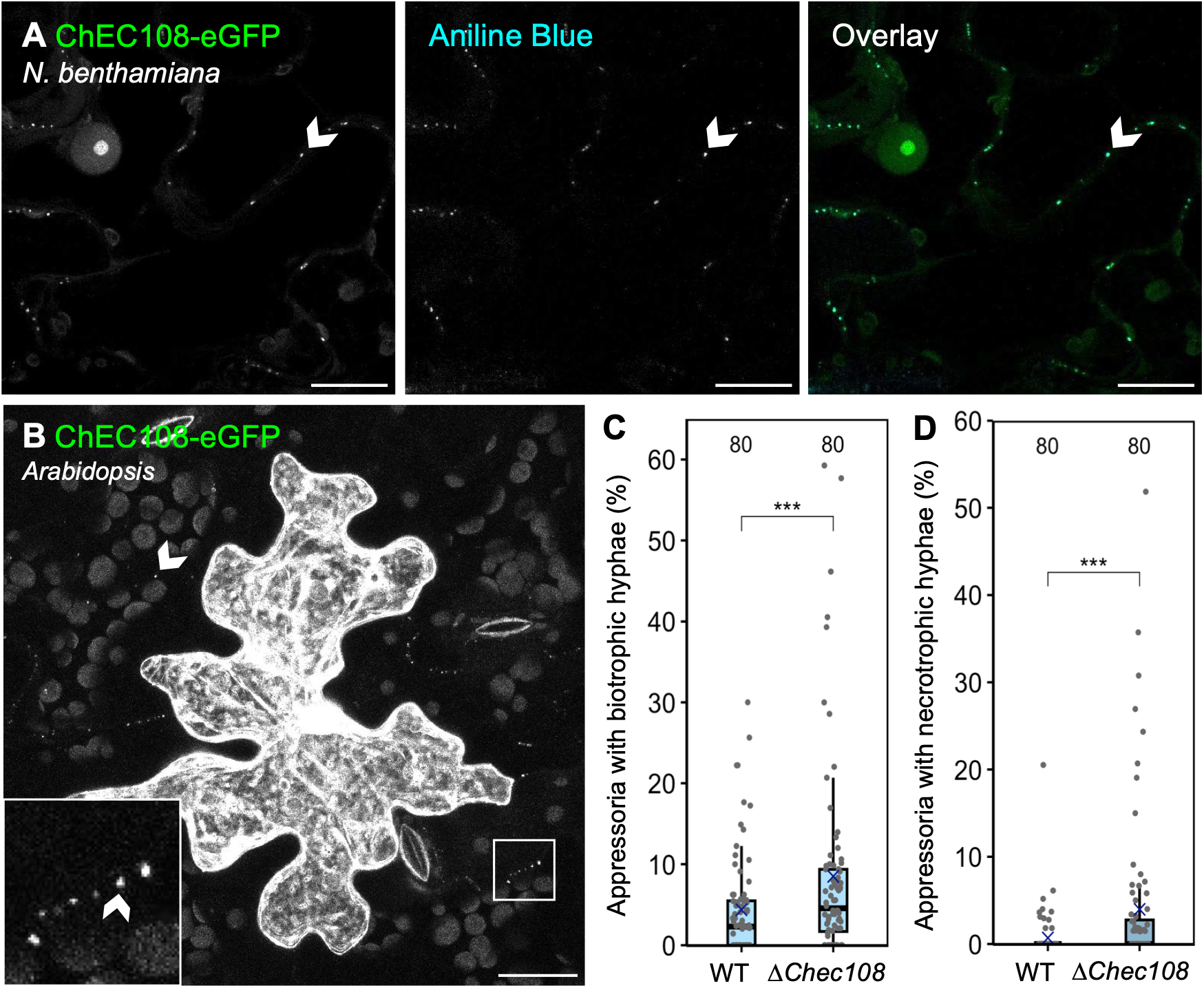
ChEC108 targets plasmodesmata and limits *C. higginsianum* virulence. **A.** *35S_pro_:ChEC108-eGFP* transiently expressed in *N. benthamiana* leaf epidermal cells via agroinfiltration, revealing plasmodesmal and nuclear localisation. ChEC108-eGFP co-localises with aniline blue-stained callose deposits. **B.** *35S_pro_:ChEC108-eGFP* transiently expressed in an Arabidopsis leaf epidermal cell via particle bombardment. ChEC108-eGFP signal can be seen at the plasmodesmata of adjacent cells, alongside chlorophyll autofluorescence. Inset panel shows a magnified view of the area indicated by a white box. Arrowheads in A and B indicate plasmodesmata. Scale bar = 20 µm. **C-D.** Percentage of appressoria associated with trypan blue-stained biotrophic hyphae (C) and necrotrophic hyphae (D) in Col-0 Arabidopsis leaves 4 days post-inoculation with wild type (WT) or *ΔChec108 C. higginsianum.* Numeric annotations indicate sample size (images), collected from 10 leaves, with 8 images captured across 3-4 inoculated zones per genotype per leaf. *** indicates p < 0.001, determined by a binomial general linear mixed model with genotype as a fixed factor, alongside leaf and zone as nested random factors.

To confirm that plasmodesmal localisation of ChEC108 is evident in a native host of *C. higginsianum*, we transiently expressed *35S_pro_:ChEC108-eGFP* in Arabidopsis leaf epidermal cells via particle bombardment. In transformed cells, fluorescent signal was observed in the nucleus and cytoplasm (Fig. 1B). Additionally, when image brightness was enhanced post-acquisition, ChEC108-eGFP could be seen labelling the plasmodesmata of adjacent cells (Fig. 1B), indicating the fusion protein targets plasmodesmata and is cell-to-cell mobile.

ChEC108-eGFP fusions accumulated to low levels in Arabidopsis cells (Fig. 1B). Therefore, we exploited the *N. benthamiana* heterologous system to explore ChEC108 function at plasmodesmata, using low optical density (OD)-agroinfiltration to achieve single-cell transformation events from which cell-to-cell mobility could be tracked. In this system, large fluorescent proteins such as 2×GFP (55 kDa) and NLS-tdTomato (62 kDa, acting as a transformation marker) typically exhibit limited cell-to-cell movement ((9), *SI Appendix,* S2, Dataset S1). By contrast, ChEC108-2×GFP (77 kDa) was highly mobile and movement of NLS-tdTomato was enhanced upon co-expression with ChEC108-2×GFP (*SI Appendix,* Fig. S2, Dataset S1). This not only supports the identification of ChEC108 as a plasmodesmata-targeting, cell-to-cell mobile *C. higginsianum* effector, but also suggests it can modulate plasmodesmal function to increase molecular flux between cells.

### ChEC108 limits *C. higginsianum* infection

In general, effectors have been proposed to function to promote pathogen virulence. To test the role of ChEC108 in *C. higginsianum* infection of Arabidopsis, we therefore generated a *C. higginsianum* null mutant lacking the effector coding sequence, referred to as *ΔChec108* (*SI Appendix,* Fig. S3). Using the *ΔChku80* background (26) as the wild-type (WT) effector control alongside the *ΔChec108* mutant, we performed virulence assays by drop-inoculating detached Arabidopsis leaves. Across repeated experiments, *ΔChec108* infection produced larger necrotrophic disease lesions in comparison to the *ΔChku80* background strain (*SI Appendix,* Fig. S4A, Dataset S2). This suggests ChEC108 has a negative influence on *C. higginsianum* infection success on Arabidopsis.

Given the narrow temporal window for effector secretion during localised biotrophy, we hypothesised that ChEC108 function could be further examined by focussing on early stages of *C. higginsianum* of infection, up to and including the transition to necrotrophic growth. We thus used trypan blue to visualise hyphal development within infected zones at the microscopic level (*SI Appendix,* Fig. S4 B and C). Relative to WT, a greater proportion of *ΔChec108* appressoria were associated with either biotrophic or necrotrophic hyphae – a trend observed both 4 and 5 days post-inoculation (Fig. 1 C and D, and *SI Appendix,* Fig. S4 D and E, Dataset S3), suggesting infection develops faster in the absence of ChEC108. To confirm that the faster rate of infection was caused by loss of ChEC108 we generated a complemented mutant fungal strain *ΔChec108.ChEC108* and assayed for biotrophic and necrotrophic hyphae 4 days post inoculation. This revealed partial complementation, with *ΔChec108.ChEC108* producing a comparable number of necrotrophic hyphae to WT, while *ΔChec108* produced more (*SI Appendix,* Fig. S5, Dataset S4). Overall, our data suggest that contrary to the expectation that effectors promote host susceptibility, ChEC108 hinders *C. higginsianum* infection of Arabidopsis.

### ChEC108 associates with HIPP6 at plasmodesmata

To investigate the mechanism by which ChEC108 targets plasmodesmata and/or activates defence, we searched for host targets of the effector. Using immunoprecipitation-mass spectrometry (IP-MS) to identify in vivo interactors in an unbiased manner, we produced ChEC108-eGFP (or eGFP alone as a control) in *N. benthamiana* leaves and examined co-immunoprecipitated proteins by MS (Dataset S5). We identified Arabidopsis homologs for the most likely ChEC108-associated proteins and sought to validate their interactions using targeted co-IP, producing ChEC108-eGFP fusions alongside mCherry-tagged candidate interactors in *N. benthamiana* leaves. Following α-GFP IP, co-enrichment of HEAVY METAL-ASSOCIATED ISOPRENYLATED PLANT PROTEIN 6 (HIPP6)-mCherry was reproducibly observed (Fig. 2A and *SI Appendix,* Fig. S6A), identifying HIPP6 (AT5G03380) as a likely binding partner of the ChEC108 effector.

**Fig. 2.**
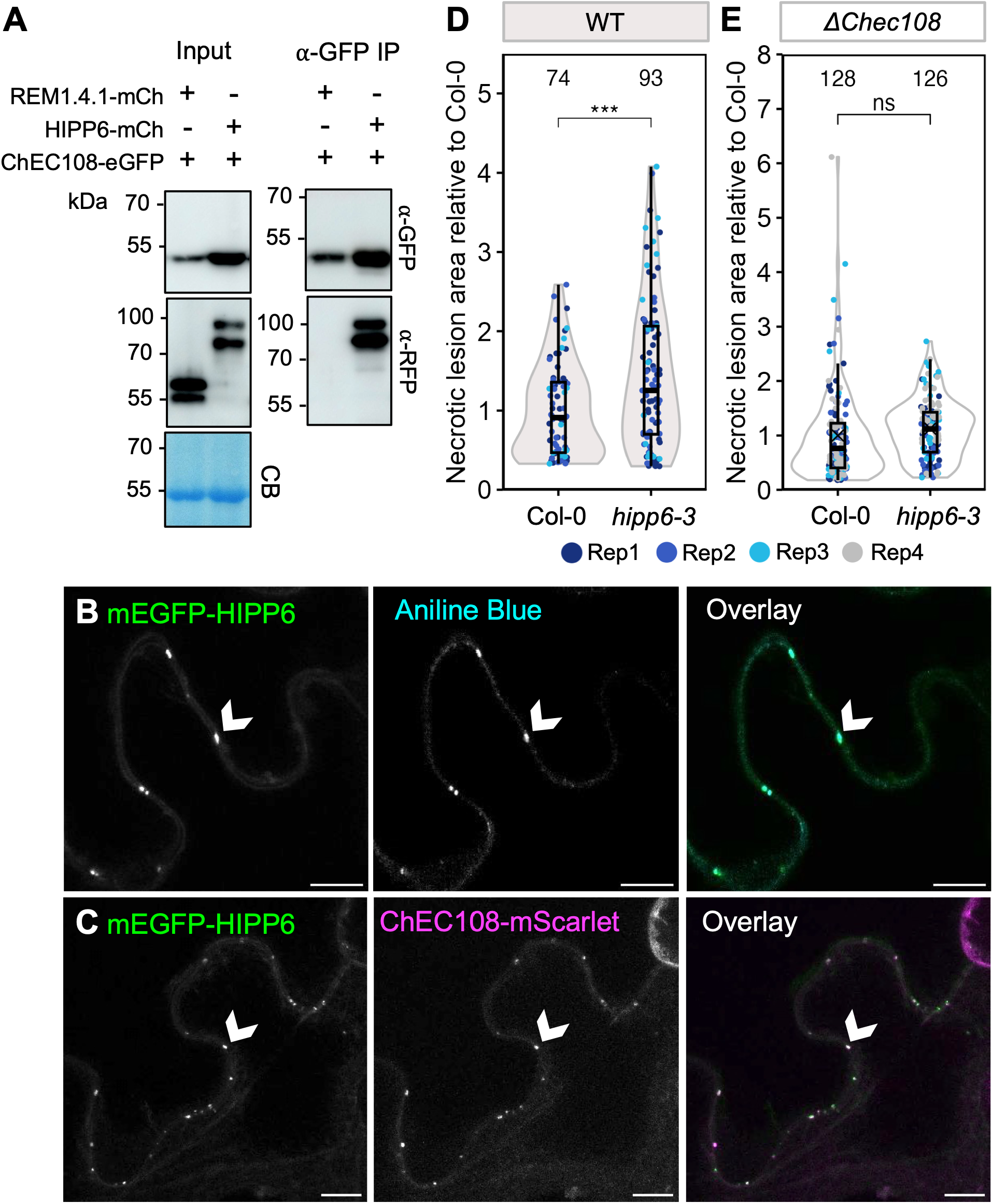
ChEC108 associates with HIPP6 at plasmodesmata. **A.** Co-IP demonstrating association of ChEC108-eGFP with mCherry (mCh)-tagged HIPP6. REM1.4.1-mCh served as a negative control. CB = Coomassie Blue staining of the Rubisco large subunit. Constructs were transiently produced in *N. benthamiana* leaves using a *35S_pro_*. Representative of three independent replicates. **B-C.** *AtUBQ10_pro_:mEGFP-HIPP6* stably expressed in Arabidopsis leaf epidermal cells. mEGFP signal is visible at puncta, which co-localise with aniline blue-stained callose deposits (B) and ChEC108-mScarlet, introduced via bombardment (C). Arrowheads indicate plasmodesmata. Scale bar = 10 µm. **D-E.** Area of necrotic lesions induced by WT (D) or *ΔChec108* (E) *C. higginsianum* on Col-0 and *hipp6-3* Arabidopsis leaves. Data were normalised by dividing all measurements by the mean area of lesions on Col-0 for a given replicate (Rep). Numeric annotations indicate sample size (lesions). *** indicates p < 0.001 and ns indicates no significant difference, determined using a linear mixed model, with replicate as a random effect and plant genotype as a fixed effect.

HIPP6 has two HMA domains (referred to as HMA1 and HMA2), and a 4-amino acid isoprenylation motif at the C-terminus. To investigate the subcellular localisation of HIPP6 in *N. benthamiana* and Arabidopsis, under the premise that a C-terminal epitope tag could block isoprenylation and induce mis-localisation, we fused fluorophores to the N-terminus. In both systems, HIPP6 fusions labelled membrane-associated puncta (*SI Appendix,* Fig. S7 A and B), which co-localised with aniline blue-stained plasmodesmal callose deposits, indicative of plasmodesmal localisation (Fig. 2B and *SI Appendix,* Fig. S7C). To confirm that ChEC108 and HIPP6 co-locate at plasmodesmata, we co-expressed *35S_pro_:ChEC108-eGFP* and *35S_pro_:mCherry-HIPP6* in *N. benthamiana*, and bombarded *35S_pro_:ChEC108-mScarlet* into leaves of Arabidopsis plants constitutively expressing *AtUBQ10_pro_:mEGFP-HIPP6*. In both scenarios, the proteins co-localised at membrane-associated puncta (Fig. 2C and *SI Appendix,* Fig. S7D). Therefore, we conclude that HIPP6 associates with ChEC108 at plasmodesmata.

To investigate the role of HIPP6 in infection, we generated a novel *hipp6-3* Arabidopsis mutant in the Col-0 background, in which the *HIPP6* coding sequence was removed by CRISPR-Cas9 genome editing (*SI Appendix,* Fig. S8, Extended Methods). We next assayed for *C. higginsianum* susceptibility by measuring the size of disease lesions that developed on *hipp6-3* mutant leaves following inoculation with the WT or *ΔChec108* fungal strain. These experiments revealed that the *hipp6-3* mutant is more susceptible than Col-0 to the WT fungus (Fig. 2D) but not to the *ΔChec108* mutant (Fig. 2E, Dataset S6). These data indicate that HIPP6 confers increased resistance to *C. higginsianum* in a ChEC108-dependent manner.

### ChEC108 binds the HMA domains of HIPP6 via metal ion co-ordination

To define the molecular interfaces underpinning ChEC108 and HIPP6 interaction, and enable targeted mutagenesis for further study, we sought structural information at atomic resolution. Prior to recombinant protein production, we identified the regions of HIPP6 required for ChEC108 binding using co-IP. As HMA domains have been previously identified as direct targets of other fungal effectors, we hypothesised that ChEC108 might associate with these regions of HIPP6 and produced ChEC108-4×Myc alongside HMA1 (amino acids 22-98 from HIPP6) and HMA2 (amino acids 152-223 from HIPP6) in *N. benthamiana*. In addition, a tandem HMA1/HMA2 fragment was produced, with the two domains connected by their native linker (referred to as tHMA, Fig. 3A). To promote protein stability of the HMA domains, a 3×FLAG-TurboID tag was fused to the N-terminus. Following co-IP, HMA1, HMA2 and tHMA were visible in the α-Myc IP fraction (*SI Appendix,* Fig. S6B), indicating that the isolated HMA domains of HIPP6 are each individually sufficient for ChEC108 association.

**Fig. 3.**
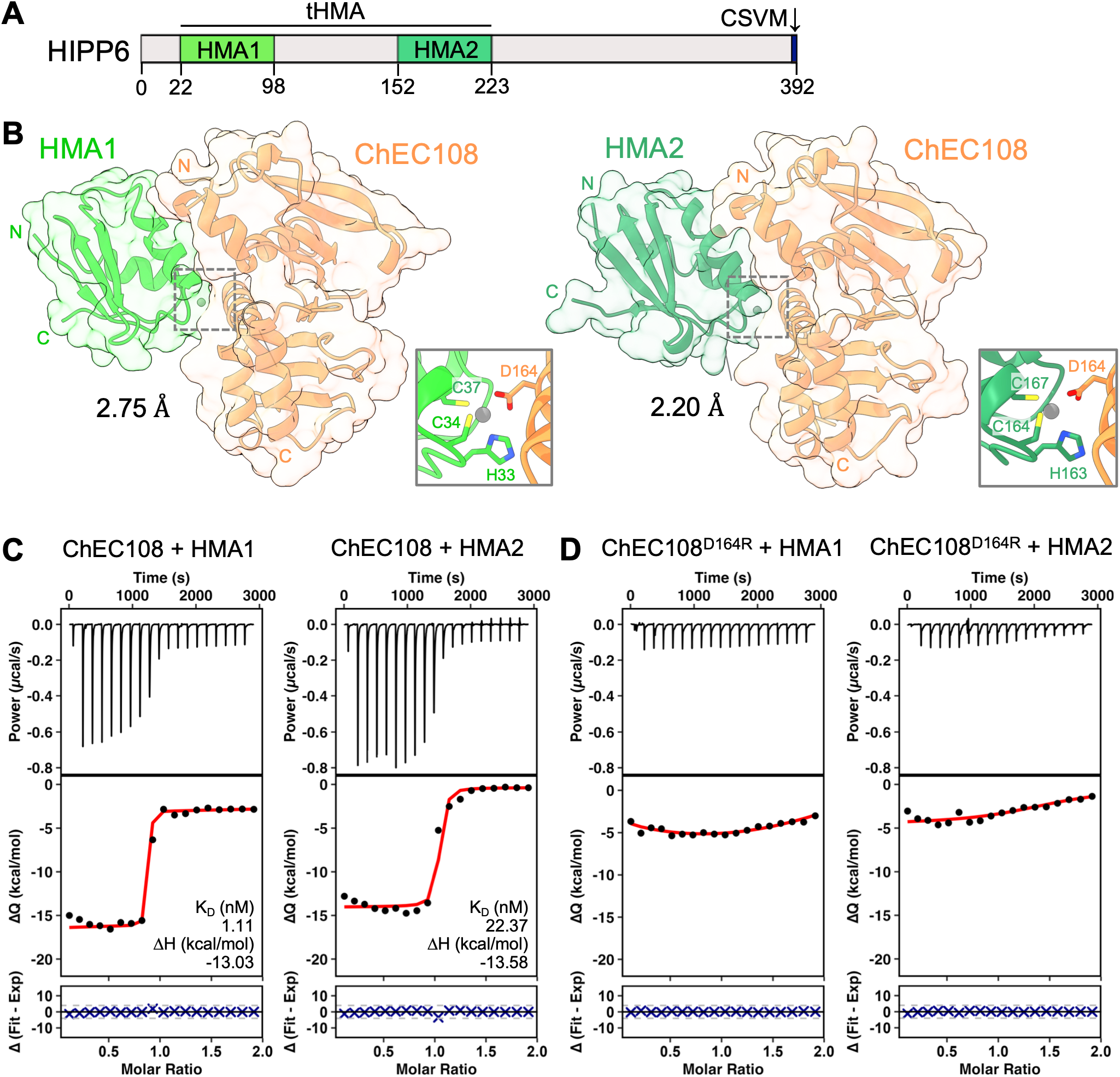
ChEC108-HMA1 and ChEC108-HMA2 complexes reveal a common binding interface involving metal ion coordination. **A.** Domain structure of HIPP6, showing HMA1, HMA2 and tandem HMA (tHMA) domain boundaries and C-terminal isoprenylation motif. **B.** Crystal structures of the ChEC108-HMA1 (PDB: 30KX) and ChEC108-HMA2 (PDB: 30KY) complexes. Grey boxes show close-up views of metal co-ordination sites at the binding interfaces. **C.** In vitro characterisation of ChEC108-HMA1 (left) or ChEC108-HMA2 binding (right) by ITC. **D.** Binding of ChEC108^D164R^ to HMA1 (left) or HMA2 (right) was not detected by ITC. In C-D, each trace represents one independent replicate, with an additional two replicates displayed in *SI Appendix*, Fig. S10.

Based on this *in planta* interaction, ChEC108-HMA1 and ChEC108-HMA2 complexes were purified from *Escherichia coli* to homogeneity, and we obtained crystals of each, refining the resulting structures to 2.75 Å and 2.20 Å resolution, respectively (*SI Appendix,* Tables S1 and S2, Extended Methods). ChEC108-HMA1 and ChEC108-HMA2 formed highly similar 1:1 complexes (RMSD = 0.795 Å across 235 pruned atom pairs, or 1.025 Å across all 244 pairs), suggesting ChEC108 adopts the same conformation when bound to either HMA domain (Fig. 3B).

ChEC108 adopts an α/β fold comprising a central eight-stranded β-sheet with nine α-helices folded against one face. This differs from the six-stranded β-sandwich fold characteristic of MAX effectors (27), which have been previously characterised in complex with HMA domains (12, 13, 17, 21). During structure refinement, a metal ion was identified at equivalent positions within the ChEC108-HMA1 and ChEC108-HMA2 binding interfaces, coordinated by three HMA-derived residues of the metal-binding β1-α1 loop (His33, Cys34 and Cys37 from HMA1, or His163, Cys164 and Cys167 from HMA2) and one ChEC108-derived residue (Asp164), in a tetrahedral configuration (Fig. 3B). X-ray fluorescence scanning of ChEC108-HMA2 complex crystals identified the associated metal as Zn^2+^. To further explore complex formation, we used analytical size exclusion chromatography (aSEC) and isothermal titration calorimetry (ITC) to study ChEC108-HMA1 and ChEC108-HMA2 interaction in vitro. In the presence of excess Zn^2+^, direct ChEC108-HMA1 and ChEC108-HMA2 binding were observed using both aSEC (*SI Appendix,* Fig. S9) and ITC, with the latter revealing ChEC108 to have nanomolar affinity for HMA1 and HMA2 (Fig. 3C and *SI Appendix,* Fig. S10 A and B).

To validate the interfaces observed in the ChEC108-HMA1 and ChEC108-HMA2 crystal structures, we designed a mutant to introduce a steric clash at the binding interface, ChEC108^D164R^. Neither ChEC108^D164R^-HMA1 nor ChEC108^D164R^-HMA2 binding could be detected by ITC (Fig. 3D and *SI Appendix,* Fig. S10 C and D). To recapitulate these findings in the context of full-length HIPP6 binding in vivo, we performed co-IP in *N. benthamiana* and observed that the ChEC108-HIPP6 association was abolished upon D164R mutation of ChEC108, or simultaneous CXXC to GXXG mutation in both HMA domains of HIPP6 (HIPP6^MM^) (*SI Appendix,* Fig. S11). Furthermore, complementation of the *ΔChec108* fungal strain with *ChEC108^D164R^* did not revert the mutant phenotype. Instead, *ΔChec108.ChEC108^D164R^* produced a greater number of necrotrophic hyphae 4 days post-inoculation than the WT fungus, comparable to the *ΔChec108* strain (*SI Appendix,* Fig. S5, Dataset S4). Thus, Asp164 from ChEC108 and the CXXC motifs of HIPP6 are critical for interaction and in vivo function via metal ion co-ordination.

In Arabidopsis, the HIPP family contains at least 45 members (11). To explore whether ChEC108 can bind the HMA domains of other HIPP family members, we selected domains from HIPPs in different phylogenetic clusters as determined by de Abreu-Neto et al. (11): cluster I (HIPP3 and HIPP5, two HMA domains), cluster II (HIPP27 and HIPP45, one HMA domain) and cluster III (HIPP34, one HMA domain). We used ITC to determine the affinity of these HMA domains for ChEC108 and ChEC108^D164R^ and observed that all HMA domains could bind ChEC108, except for HMA2 from HIPP5 and the HIPP27 HMA domain (*SI Appendix,* Table S3 and Fig. S12). No HMA domains tested could bind ChEC108^D164R^, while the HMA domains of HIPP6 bound ChEC108 with the highest affinity (*SI Appendix,* Table S3). Therefore, we conclude that ChEC108 can bind other HIPPs via metal ion co-ordination at the same interface, raising the possibility of functional redundancy between HIPPs in this context.

### ChEC108 plasmodesmal localisation and mobility do not require HIPP6 binding

Equipped with structure-guided mutants, we sought to determine whether HIPP6 binding is required for ChEC108 localisation and mobility. We generated transgenic Arabidopsis lines that express *AtUBQ10_pro_:ChEC108-eGFP* or *AtUBQ10_pro_:ChEC108^D164R^-eGFP* and observed that in two independent transgenic lines for each construct, both ChEC108-eGFP and ChEC108^D164R^-eGFP were observed exclusively at plasmodesmata (*SI Appendix,* Fig. S13A), suggesting ChEC108 plasmodesmal localisation does not depend on HIPP6. To investigate the capacity of ChEC108 to move through plasmodesmata when unable to bind HIPP6, we used particle bombardment to introduce ChEC108-eGFP into single cells of the *hipp6-3* Arabidopsis mutant and observed that the effector remained cell-to-cell mobile (Fig. 4A). Likewise, the non-HMA binding variant ChEC108^D164R^-eGFP was able to move cell-to-cell in Col-0 leaves (Fig. 4B). This indicates that HIPP binding is not necessary for ChEC108 movement and that ChEC108 must rely on other, currently unknown factors to direct it towards and/or through plasmodesmata in host cells.

**Fig. 4.**
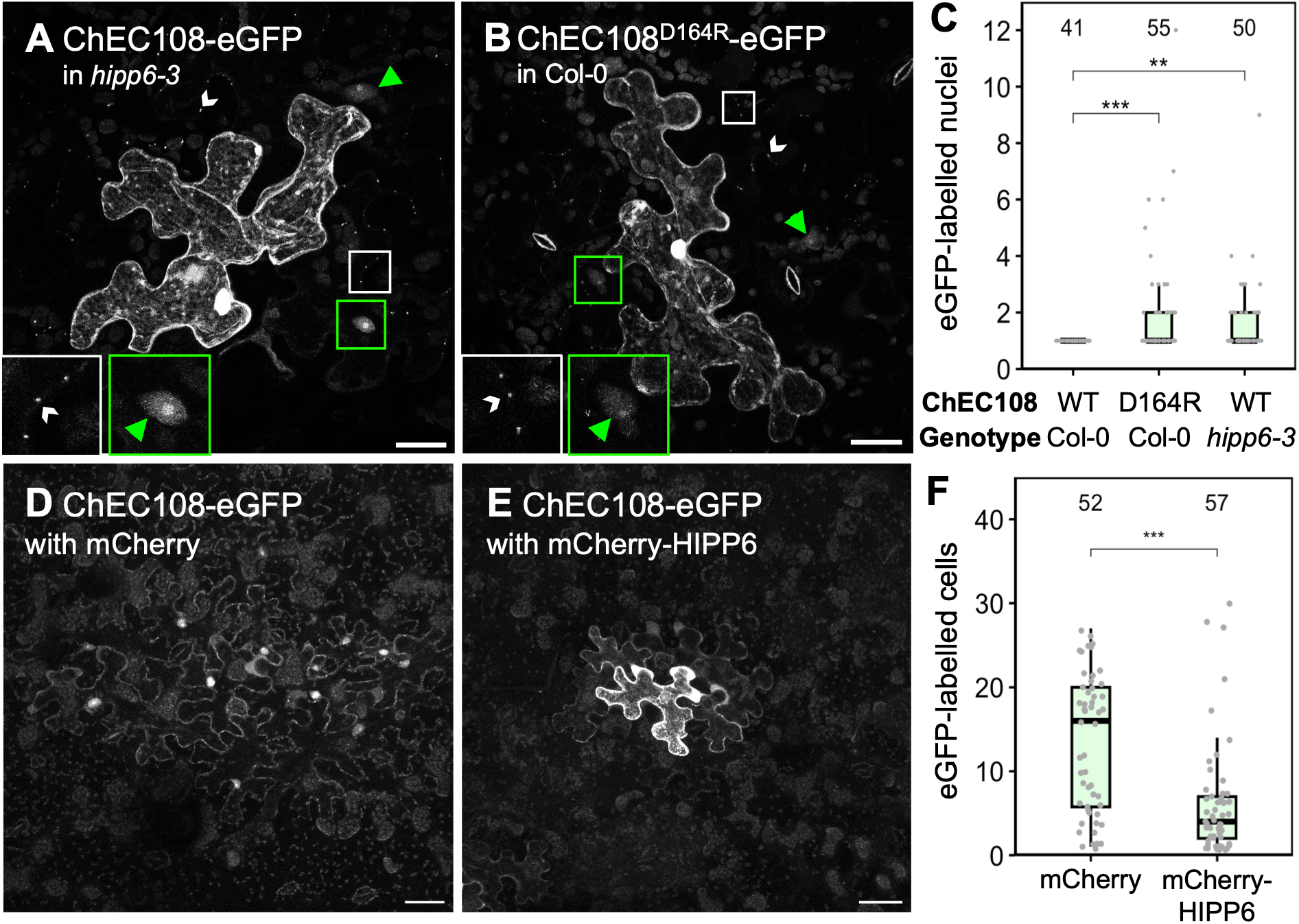
Cell-to-cell mobility of ChEC108 is reduced upon ChEC108-HIPP6 association. A-B. *35S_pro_:ChEC108-eGFP* (A) or *35S_pro_:ChEC108^D164R^-eGFP* (B) transiently expressed via bombardment in an Arabidopsis leaf epidermal cell of genotype *hipp6-3* (A) or Col-0 (B). eGFP signal is visible at plasmodesmata (white arrowheads) in adjacent cells, alongside chlorophyll autofluorescence. Green arrowheads indicate nuclei of adjacent cells. Inset panels show a magnified view of areas indicated by white and green boxes. Scale bar = 20 µm. **C.** Number of nuclei labelled by ChEC108-eGFP or ChEC108^D164R^-eGFP at bombardment sites in Col-0 or *hipp6-3* leaves. Numeric annotations indicate sample size (sites), collected across 5-6 leaves per treatment, with a maximum of 14 sites per leaf. *** indicates p < 0.001 and ** indicates p < 0.01, determined by Kruskal-Wallis and post-hoc Dunn tests. **D-E.** *35S_pro_:ChEC108-eGFP* transiently expressed in single *N. benthamiana* epidermal cells using low-OD agroinfiltration of leaves transiently expressing *35S_pro_:mCherry* (D) or *35S_pro_:mCherry-HIPP6* (E) using high-OD agroinfiltration. Scale bar = 50 µm. **F.** Mobility of ChEC108-eGFP quantified 2 days post-infiltration by counting the number of fluorescent cells at transformation sites. Numeric annotations indicate sample size (sites), collected across 3-4 leaves per treatment, with a maximum of 20 sites per leaf. *** indicates p < 0.001, determined by bootstrapping analysis with 5000 iterations.

### ChEC108 binding to HIPP6 induces plasmodesmal closure

Close examination of ChEC108^D164R^-eGFP localisation at bombardment sites in Col-0, and of ChEC108-eGFP at bombardment sites in *hipp6-3* leaves suggested that relative to ChEC108-eGFP in Col-0 leaves (Fig. 1B), these contexts show an elevated nuclear signal in the adjacent cells (green arrows in Fig. 4 A and B). Quantification of this phenomenon revealed significantly more nuclei were visible in cells around those transformed in scenarios where ChEC108-HIPP6 binding was compromised (that is, ChEC108^D164R^-eGFP in Col-0, or ChEC108-eGFP in *hipp6-3*), in comparison to when ChEC108-HIPP6 binding was possible (ChEC108-eGFP in Col-0) (Fig. 4C, Dataset S7). This finding supports a model whereby HIPP6 binding ordinarily inhibits ChEC108 movement in host leaves.

ChEC108 is highly mobile in *N. benthamiana* (a non-host of *C. higginsianum*) (*SI Appendix,* Fig. S2) yet our IP-MS data suggest the effector can bind an *N. benthamiana* ortholog of HIPP6 (*SI Appendix,* Dataset S5), raising the possibility that HIPP6 might behave differently in these two species. To explore this, we conducted a mobility assay in *N. benthamiana* leaves in the presence and absence of Arabidopsis HIPP6. Using low OD-agroinfiltration to express *35S_pro_:ChEC108-eGFP* at discrete single cell sites, we measured the spread of eGFP signal in leaves co-transformed with *35S_pro_:mCherry-HIPP6*, or *35S_pro_:mCherry* as a negative control (Fig. 4 D and E). Consistent with our previous observations (*SI Appendix,* Fig. S2 and Dataset S1), ChEC108-eGFP proved highly mobile in control leaves expressing *mCherry,* although its mobility was restricted when co-expressed with *mCherry-HIPP6* (Fig. 4F and *SI Appendix,* Dataset S8). This supports the hypothesis that Arabidopsis HIPP6 inhibits ChEC108 movement via plasmodesmata.

To test whether ChEC108 binding to HIPP6 has consequences for global cell-to-cell trafficking beyond restriction of ChEC108 movement, we generated transgenic Arabidopsis lines constitutively expressing *AtUBQ10_pro_:ChEC108-eGFP* or *AtUBQ10_pro_:ChEC108^D164R^-eGFP*. Relative to Col-0 plants, those expressing *ChEC108-eGFP* were severely dwarfed and developed small, curled leaves, while those expressing *ChEC108^D164R^-eGFP* were comparable to Col-0 (*SI Appendix,* Fig. 14A). We assessed baseline plasmodesmal aperture by transiently expressing *35S_pro_:mScarlet* via particle bombardment and observed extensive spread of mScarlet signal in Col-0 and *ChEC108^D164R^-eGFP*-expressing leaves. However, in leaves expressing *ChEC108-eGFP*, mScarlet was severely restricted, often confined to single cells (*SI Appendix,* Fig. S14 B and C, Dataset S9). Consistent with the growth phenotypes observed (*SI Appendix,* Fig. S14A), this suggests ChEC108 binding to HIPP6 promotes plasmodesmal closure, compromising Arabidopsis cell-to-cell communication and affecting multiple cellular processes.

### ChEC108 induces defence-related gene expression in Arabidopsis

To explore the processes perturbed by ChEC108-HIPP6 binding, we compared transcriptomes of Col-0, *ChEC108-eGFP*-and *ChEC108^D164R^-eGFP*-expressing Arabidopsis seedlings, and identified differentially expressed genes (DEGs, defined as showing log_2_ fold change ≥ |1| and Wald test adjusted p value ≤ 0.05) for pairwise genotype comparisons (*SI Appendix,* Dataset S10). This revealed that *ChEC108-eGFP-*expressing seedlings displayed a marked shift in their transcriptomic profile, with 391 upregulated and 48 downregulated genes versus Col-0, while seedlings expressing *ChEC108^D164R^-eGFP* were much less perturbed, with 21 upregulated and 5 downregulated genes compared to Col-0. Focussing on upregulated processes, we assessed the overlap of DEGs between the three genotypes and identified genes induced only in the presence of ChEC108-eGFP (393 genes, *SI Appendix,* Fig. S15A). Within this list were genes associated with defence-and immune-related GO terms, including ‘response to salicylic acid’, ‘systemic acquired resistance’ and ‘plant-type hypersensitive response’ (*SI Appendix,* Fig. S15B, Dataset S11). Given that few transcriptional changes were observed in the presence of ChEC108^D164R^-eGFP, these data support a model in which ChEC108-HIPP6 interaction induces plasmodesmal closure and stress responses in Arabidopsis, culminating in growth inhibition.

## Discussion

In plants, HMA domain-containing proteins have frequently emerged as binding partners of pathogen effectors, but the intracellular consequences of these interactions are not well-understood. Here, we report that binding of a plasmodesmata-targeting effector, ChEC108, to the Arabidopsis plasmodesmal HMA protein, HIPP6, triggers defence responses. Bringing together results from imaging-based mobility assays, transcriptomic analysis and fungal virulence assays, we propose that HIPP6 binding of ChEC108 at plasmodesmata initiates plasmodesmal closure, thereby restricting cell-to-cell movement of the effector and initiating host defence signalling (*SI Appendix,* Fig. S16). This impairs *C. higginsianum* infection, suggesting that HIPP6 functions in a plasmodesmata-located surveillance system that activates immune responses upon pathogen detection.

HIPP proteins from multiple species have been identified as plasmodesmal components via live imaging studies (15, 16, 18, 28) and proteomic analysis of Arabidopsis, *N. benthamiana* and *Populus trichocarpa* subcellular fractions (29–34). However, with HIPP6 orthologs present in both scenarios, our mobility assays in *N. benthamiana* (in which ChEC108 is highly mobile) and Arabidopsis leaves (in which ChEC108 mobility was low) suggest functional differences exist even between HIPP6 orthologs. Indeed, in other systems, HMA proteins have been identified as both susceptibility (15, 35–37) and resistance-promoting (38, 39) factors. Here, our data suggest Arabidopsis HIPP6 functions as an activator of immunity and a positive regulator of resistance.

As an evolutionarily maintained effector, it is reasonable to hypothesise that ChEC108 fulfills a virulence function that benefits the reproductive success of the fungus. However, our data suggest that any virulence function is masked in the context of Arabidopsis infection. The Pwl2 effector from *M. oryzae* binds plasmodesmata-located HIPP43 orthologs in rice and barley (13, 16), as well as the barley NLR MLA3 (40), which carries a structural mimic of HIPP43 at its C-terminus (41). A *Δpwl2 M. oryzae* mutant is more virulent on weeping lovegrass and *+Mla3* barley cultivars, but less virulent on rice (16), meaning Pwl2 differentially benefits *M. oryzae* on different hosts. Our observation that HIPP6 ortholog function varies in different species allows for the hypothesis that ChEC108 has been evolutionarily maintained because it has an unimpeded virulence function in other *C. higginsianum* host species.

An alternative hypothesis to explain the evolutionary maintenance of ChEC108, might be that stress responses induced by the effector could potentially contribute to *C. higginsianum* fitness if properly temporally controlled, for example by fulfilling a function required for transition from biotrophy to necrotrophy. Consistent with this idea, stage-specific transcriptomic data analysed by Dallery et al. (42) suggest that *ChEC108* expression peaks during the biotrophic stage of infection on Arabidopsis. However, as the *ΔChec108* mutant exhibited faster progression to biotrophic and necrotrophic growth, as well as greater disease development, our data contradict the idea that ChEC108 is a driver of necrotrophy.

We observed that *ChEC108*-expressing leaves, but not *ChEC108^D164R^*-expressing leaves, have reduced cell-to-cell connectivity, suggesting that ChEC108-HIPP6 binding is associated with plasmodesmal closure. Restriction of plasmodesmata is an established immune response that occurs downstream of bacterial (flg22) and fungal (chitin) elicitor perception (1, 3). Further, Tee et al. (43) demonstrated that this response induces stress, involving accumulation of the defence hormone salicylic acid, as well as transcriptional induction of defence genes. Given that plasmodesmal closure and immune-related gene expression were enhanced upon *ChEC108* expression in Arabidopsis, it is possible that effector binding perturbs HIPP6-dependent regulation of plasmodesmal aperture, thereby initiating downstream defence signalling.

Effectors from diverse pathogens have been found to target plasmodesmata, identifying them as a hub for pathogen invasion. For example, *Pseudomonas syringae* pv. *tomato* (*Pst*) DC3000 (44), *Fusarium oxysporum* (45, 46), and viruses (47, 48) target plasmodesmata to enhance movement of pathogen molecules between host cells and thus promote virulence. In this context, a plasmodesmata-located surveillance and defence mechanism would offer an additional means by which the host could impede pathogen invasion and might explain how HIPP6 has evolved such a defensive function.

Our study has therefore revealed that an Arabidopsis HMA protein, HIPP6, functions in plasmodesmal immune surveillance and that when infecting Arabidopsis, *C. higginsianum* produces an effector, ChEC108, that binds HIPP6, thereby activating plasmodesmal closure and downstream immune responses, impeding infection. A plasmodesmal effector-triggered immune response therefore impedes cell-to-cell trafficking during fungal invasive growth.

## Materials and Methods

Extended Methods are available in the *SI Appendix*.

### Genetic material

The Δ*Chec108* strain of *C. higginsianum* was generated in the *ΔChku80* background (26) which is optimised for homologous recombination. We used targeted homologous recombination to replace the coding sequence of *ChEC108* with a hygromycin resistance cassette, achieved by agrobacterium-mediated transformation of fungal conidia. Complemented lines were generated in the Δ*Chec108* background by non-targeted insertion of a *ChEC108_pro_:ChEC108:ChEC108_ter_* or *ChEC108_pro_:ChEC108^D164R^:ChEC108_ter_* expression cassette, accompanied by a Basta resistance marker. Arabidopsis *hipp6-3* mutants were generated in the Columbia (Col-0) background, using CRISPR-Cas9 to fully excise the *HIPP6* coding sequence (*SI Appendix,* Extended Methods). Arabidopsis overexpression lines were generated in the Col-0 background by agrobacterium-mediated insertion of *AtUBQ10_pro_::ChEC108-eGFP::t35S_ter_* or *AtUBQ10_pro_::ChEC108^D164R^-eGFP::t35S_ter_* cassettes, alongside a kanamycin resistance marker.

### Agroinfiltration of Nicotiana benthamiana

Plasmids were transformed into *Agrobacterium tumefaciens* GV3101 by electroporation, and cultures were diluted in infiltration buffer (10 mM MES pH 5.7, 10 mM MgCl_2_, 100 µM acetosyringone) prior to infiltration of mature leaves on 3-4-week-old *N. benthamiana* plants. To achieve widespread transformation events across a leaf, bacteria were diluted to an OD_600_ of 0.1-0.4. To achieve discrete single cell transformation events for quantification of protein mobility, bacteria were diluted to an OD_600_ of 0.0001. All infiltration mixtures were prepared with bacteria carrying the RNA silencing suppressor p19 at OD_600_ = 0.1. Leaf material was harvested 2-3 days post infiltration.

Protein subcellular localisation and cell-to-cell movement in *N. benthamiana* was analysed using an LSM Zeiss 800 confocal microscope, equipped with a 63× (W Plan-Apochromat 63×/1.0 M27, Zeiss) or 20× (W N-Achroplan 20×/0.5 M27, Zeiss) water-dipping objective. To visualise callose deposits at plasmodesmata, leaves were first infiltrated with 0.01% aniline blue and imaged using a 63× (C-Apochromat 63×/1.2 W Corr M27, Zeiss) water-immersion objective.

### Bombardment of Arabidopsis

Microprojectile bombardment of fully expanded Arabidopsis leaves, detached from 5-6-week-old rosettes, was carried out as described by Tee et al. (49) using a Biolistic PDS-1000/He particle delivery system (BIO-RAD). After 18-24h, bombarded leaves were imaged using an LSM Zeiss 800 confocal microscope, equipped with a 63× (W Plan-Apochromat 63×/1.0 M27, Zeiss) water-dipping objective. Nuclear counts were performed manually.

### Colletotrichum higginsianum infection of Arabidopsis

Fungal colonies were cultured for 2-3 weeks in the dark on Mathur’s Medium. Conidial suspensions were collected in water and diluted to 2×10^6^ conidia/mL (for macroscopic lesion measurements) or 2×10^5^ conidia/mL (for microscopic study of infection structures) and used to drop-inoculate the adaxial surface of leaves detached from 5-6-week-old Arabidopsis plants. Leaves were sealed with parafilm inside 1.5% agar plates and incubated at 22 °C with a 10 h light/14 h dark photoperiod. For macroscopic lesion measurements, one 3 µl droplet was applied to each leaf, and lesions were photographed 6-8 days post-inoculation.

For microscopic study of infection structures, four 1 µL droplets were applied per genotype per leaf, and 6 mm leaf discs were harvested 4 and 5 days post-inoculation. Leaf discs were boiled with trypan blue staining solution [25% (v/v) glycerol, 25% (v/v) lactic acid, 25% (v/v) water-saturated phenol, 0.025% (w/v) trypan blue] at 95 °C for 2.5 min, submerged in staining solution for 30 min at room temperature, then rinsed with water and destained in a 5:2 ratio (w/v) chloral hydrate/water solution with gentle agitation. When sufficiently destained, leaf discs were mounted in 40% glycerol and imaged using a Zeiss Axio Imager Z2, equipped with a 20× (EC Plan-Neofluar 20×/0.5, Zeiss) air-immersion objective. Appressoria and hyphal counts were conducted manually.

### Co-immunoprecipitation

Proteins from *N. benthamiana* leaf tissue were extracted in IP buffer [50 mM Tris-HCl pH 7.5, 150 mM NaCl, 1 mM EDTA pH 8.0, 10% glycerol, 0.5% NP40 IGEPAL® CA-630, cOmplete^TM^ ULTRA EDTA-free protease inhibitor tablets (1 per 50 mL) (Roche), PhosSTOP EASYpack phosphatase inhibitor cocktail tablets (1 tablet per 50 mL) (Roche), 1 mM Na_2_MoO_4_×2H_2_O, 1mM NaF, 5 mM DTT and 1mM PMSF]. To enrich for GFP-or Myc-tagged proteins, extracts were mixed with 20 µl of GFP-Trap® magnetic agarose beads (Chromotek gtma-20) or 80-100 µl of Pierce™ anti-c-Myc agarose bead slurry (Thermo Scientific 20168) and rotated end-over-end for 2-5 h at 5 °C. After, beads were washed five times in IP buffer. Input and IP samples were boiled with 8-10 µl of loading buffer [100 mM Tris, pH 6.8, 42% (v/v) glycerol, 2.5% (w/v) SDS, 0.04% (w/v) bromophenol blue, 10% (v/v) β-mercaptoethanol] at 95 °C for 15 min prior to SDS-PAGE. Western blotting details can be found in the *SI Appendix,* Extended Methods.

### Protein production from *Escherichia coli*

SHuffle *E. coli* (50) cells were transformed and cultured at 30 °C until the OD_600_ reached 0.6-0.8. Expression was induced at 18 °C by adding 1 mM isopropyl β-D-1-thiogalactopyranoside (IPTG). Induction proceeded for 17-24 h before cells were pelleted by centrifugation, resuspended and lysed by sonication. The cleared lysate was loaded onto a 5 mL HisTrap^TM^ HP NTA column (Cytiva) for immobilised metal affinity chromatography (IMAC), followed by size exclusion chromatography (SEC). Affinity tags were cleaved by overnight incubation with 6×His-tagged human rhinovirus 3C protease (10 µg/mg of protein) at 4 °C. 3C protease, cleaved affinity tags and remaining uncleaved contaminants were removed by IMAC, and proteins of interest were concentrated using VivaSpin® 20 mL concentrators (Sartorius) with a 3 kDa, 10 kDa, or 30 kDa molecular weight cut-off. Samples were further purified by SEC, concentrated as above and frozen in liquid nitrogen for storage at −70 °C. For in vitro binding assays, proteins were purified according to Yu et al. (23).

### **X-** ray crystallography

Initial crystallisation trials were conducted using the sitting drop vapour diffusion method with commercially available sparse matrix screens, dispensed into MRC 96-Well 2-Drop Plates (Molecular Dimensions). ChEC108-HMA1 crystals formed in 25% Polyethylene glycol monomethyl ether (PEG-MME) 2K, 0.1 M HEPES pH 7.5 (well B6, PEGs Suite, QIAGEN) using an initial protein concentration of 20 mg/mL. ChEC108-HMA2 crystals were formed in 16% PEG 6K, 0.1 M TRIS pH 7.5, using an initial protein concentration of 20 mg/mL. Prior to X-ray diffraction, crystals were cryoprotected in ethylene glycol and vitrified in liquid nitrogen. X-ray diffraction data were collected at beamline i24 (MX32728-3 for ChEC108-HMA1) and i04 (MX32728-5 and MX32728-16 for ChEC108-HMA2) of the Diamond Light Source synchrotron facility (Oxford, UK). For details on data processing, see *SI Appendix*, Extended Methods.

### Isothermal titration calorimetry

In vitro purified proteins (ChEC108, ChEC108^D164R^, or HMA domains) were diluted in SEC buffer (20 mM HEPES, pH 7.5, 150 mM NaCl) supplemented with 1 mM DTT and 50 µM ZnCl_2_, then loaded into the experimental cell (250 µl of HMA domain) or syringe (50 µl of ChEC108 or ChEC108^D164R^) of a MicroCal PEAQ-ITC machine (Malvern, UK). For details, see *SI Appendix* Extended Methods.

## Statistical analysis

All statistical tests were conducted in R. Details to accompany each figure are given in Dataset S12.

## Data availability

Sequence data can be found in the GenBank/EMBL data libraries under accession numbers CH63R_09563 (*ChEC108*) and AT5G03380 (*HIPP6*). Crystal structures of the ChEC108-HMA1 and ChEC108-HMA2 complexes have been deposited in the Protein Data Bank under accession codes 30KX and 30KY, respectively. RNA-sequencing data has been deposited on ArrayExpress under the accession code E-MTAB-16996.

## Acknowledgments and funding sources

We acknowledge access to the John Innes Centre Bioimaging Facility and would like to thank Dr Eva Wegel for training and assistance with light microscopy experiments. We thank Prof. David Lawson and Julia Mundy of the John Innes Centre Structural Biology platform for their assistance with protein crystallisation and X-ray data collection. We thank Abass Maqbool of the John Innes Centre Biophysical Analysis platform for assistance with ITC experiments, as well as the John Innes Centre Laboratory Support and Horticultural Services teams. For funding, we acknowledge the United Kingdom Research and Innovation (UKRI) Biotechnology: Engineering and Physical Sciences Research Council (UKRI Frontier Research Guarantee Grants EP/Z534390/1 ‘ACUTE’ and EP/X022439/1 ‘SEPBLAST’) and Biological Sciences Research Council (BBSRC, grant numbers: BB/X010996/1; BB/X007685/1; BB/Y008782/1; BB/X016056/1). Work was also funded by the European Research Council (grants 725459 ‘INTERCELLAR’), the John Innes Foundation (Rotation PhD studentship) and the Gatsby Charitable Foundation.

## References

1. C. Cheval, et al., Chitin perception in plasmodesmata characterizes submembrane immune-signaling specificity in plants. Proc Natl Acad Sci U S A 117, 9621–9629 (2020).

2. C. Faulkner, et al., LYM2-dependent chitin perception limits molecular flux via plasmodesmata. Proceedings of the National Academy of Sciences 110, 9166–9170 (2013).

3. B. Xu, et al., A calmodulin-like protein regulates plasmodesmal closure during bacterial immune responses. New Phytologist 215, 77–84 (2017).

4. M. Dickmanns, et al., In situ architecture of plasmodesmata in *Physcomitrium patens* resolved by cryo-electron tomography. Nat. Plants 12, 1051–1061 (2026).

5. U. Damm, R. J. O’Connell, J. Z. Groenewald, P. W. Crous, The *Colletotrichum destructivum* species complex - hemibiotrophic pathogens of forage and field crops. Stud. Mycol. 79, 49–84 (2014).

6. R. O’Connell, et al., A Novel *Arabidopsis*-*Colletotrichum* Pathosystem for the Molecular Dissection of Plant-Fungal Interactions. Molecular Plant-Microbe Interactions 17, 272–282 (2004).

7. R. J. O’Connell, et al., Lifestyle transitions in plant pathogenic *Colletotrichum* fungi deciphered by genome and transcriptome analyses. Nat. Genet. 44, 1060–1065 (2012).

8. J. Kleemann, et al., Sequential Delivery of Host-Induced Virulence Effectors by Appressoria and Intracellular Hyphae of the Phytopathogen *Colletotrichum higginsianum*. PLoS Pathog. 8, e1002643 (2012).

9. M. Ohtsu, et al., Assaying effector cell-to-cell mobility in plant tissues identifies hypermobility and indirect manipulation of plasmodesmata. Molecular Plant-Microbe Interactions (2023). 10.1094/MPMI-05-23-0052-TA.

10. G. P. Robin, et al., Subcellular localization screening of *Colletotrichum higginsianum* effector candidates identifies fungal proteins targeted to plant peroxisomes, golgi bodies, and microtubules. Front. Plant Sci. 9 (2018).

11. J. B. De Abreu-Neto, A. C. Turchetto-Zolet, L. F. V. De Oliveira, M. H. Bodanese Zanettini, M. Margis-Pinheiro, Heavy metal-associated isoprenylated plant protein (HIPP): Characterization of a family of proteins exclusive to plants. FEBS Journal 280, 1604–1616 (2013).

12. A. R. Bentham, et al., A single amino acid polymorphism in a conserved effector of the multihost blast fungus pathogen expands host-target binding spectrum. PLoS Pathog. 17, e1009957 (2021).

13. R. Zdrzałek, et al., Bioengineering a plant NLR immune receptor with a robust binding interface toward a conserved fungal pathogen effector. Proceedings of the National Academy of Sciences 121, e2402872121 (2024).

14. J. H. R. Maidment, et al., Multiple variants of the fungal effector AVR-Pik bind the HMA domain of the rice protein OsHIPP19, providing a foundation to engineer plant defense. Journal of Biological Chemistry 296, 100371 (2021).

15. K. Oikawa, et al., The blast pathogen effector AVR-Pik binds and stabilizes rice heavy metal-associated (HMA) proteins to co-opt their function in immunity. PLoS Pathog. 20, e1012647-(2024).

16. V. M. Were, et al., The *Magnaporthe oryzae* effector Pwl2 alters HIPP43 localization to suppress host immunity. Plant Cell 37, koaf116 (2025).

17. J. H. R. Maidment, et al., Two folds, many faces: The *Magnaporthe oryzae* MAX effector AVR-Pia targets novel rice HMA domain-containing proteins. PLoS Pathog. 22, e1014382-(2026).

18. G. H. Cowan, et al., Potato Mop-Top Virus Co-Opts the Stress Sensor HIPP26 for Long-Distance Movement. Plant Physiol. 176, 2052–2070 (2018).

19. X. Niu, et al., A small secreted protein, RsMf8HN, in *Rhizoctonia solani* triggers plant immune response, which interacts with rice OsHIPP28. Microbiol. Res. 266, 127219 (2023).

20. H. Song, et al., The *Meloidogyne graminicola* effector MgMO289 targets a novel copper metallochaperone to suppress immunity in rice. J. Exp. Bot. 72, 5638–5655 (2021).

21. A. Maqbool, et al., Structural basis of pathogen recognition by an integrated HMA domain in a plant NLR immune receptor. Elife 4, e08709 (2015).

22. S. Cesari, et al., The Rice Resistance Protein Pair RGA4/RGA5 Recognizes the *Magnaporthe oryzae* Effectors AVR-Pia and AVR1-CO39 by Direct Binding. Plant Cell 25, 1463–1481 (2013).

23. D. S. Yu, et al., Engineering plant tandem kinase immune receptors expands effector recognition profiles. Sci. Adv. 12, eaef2426 (2026).

24. Z. Bernasconi, et al., An HMA-like integrated domain in the wheat tandem kinase WTK4 recognises an RNase-like pathogen effector. Nat. Commun. 17, 8458 (2026).

25. C. H. Khang, et al., Translocation of *Magnaporthe oryzae* Effectors into Rice Cells and Their Subsequent Cell-to-Cell Movement. Plant Cell 22, 1388–1403 (2010).

26. M. Korn, et al., A Genetic Screen for Pathogenicity Genes in the Hemibiotrophic Fungus *Colletotrichum higginsianum* Identifies the Plasma Membrane Proton Pump Pma2 Required for Host Penetration. PLoS One 10, e0125960 (2015).

27. K. De Guillen, et al., Structure Analysis Uncovers a Highly Diverse but Structurally Conserved Effector Family in Phytopathogenic Fungi. PLoS Pathog. 11, e1005228 (2015).

28. T. Guo, et al., Arabidopsis HIPP proteins regulate endoplasmic reticulum-associated degradation of CKX proteins and cytokinin responses. Mol. Plant 14, 1918–1934 (2021).

29. M. G. Johnston, et al., Comparative phyloproteomics identifies conserved plasmodesmal proteins. J. Exp. Bot. 74, 1821–1835 (2023).

30. L. Fernandez-Calvino, et al., Arabidopsis Plasmodesmal Proteome. PLoS One 6, e18880 (2011).

31. S.-H. Park, et al., NbEXPA1, an α-expansin, is plasmodesmata-specific and a novel host factor for potyviral infection. The Plant Journal 92, 846–861 (2017).

32. F. Leijon, M. Melzer, Q. Zhou, V. Srivastava, V. Bulone, Proteomic Analysis of Plasmodesmata From *Populus* Cell Suspension Cultures in Relation With Callose Biosynthesis. Front. Plant Sci. 9 (2018).

33. M. L. Brault, et al., Multiple C2 domains and transmembrane region proteins (MCTPs) tether membranes at plasmodesmata. EMBO Rep. 20, e47182 (2019).

34. M. E. Kraner, C. Müller, U. Sonnewald, Comparative proteomic profiling of the choline transporter-like1 (CHER1) mutant provides insights into plasmodesmata composition of fully developed *Arabidopsis thaliana* leaves. The Plant Journal 92, 696–709 (2017).

35. S. Fukuoka, et al., Loss of Function of a Proline-Containing Protein Confers Durable Disease Resistance in Rice. Science (1979). 325, 998–1001 (2009).

36. Z. S. Radakovic, et al., Arabidopsis *HIPP27* is a host susceptibility gene for the beet cyst nematode *Heterodera schachtii*. Mol. Plant Pathol. 19, 1917–1928 (2018).

37. Q. M. Imran, et al., Nitric Oxide Responsive Heavy Metal-Associated Gene *AtHMAD1* Contributes to Development and Disease Resistance in *Arabidopsis thaliana*. Front. Plant Sci. 7 (2016).

38. L.-X. Chai, et al., A putative nuclear copper chaperone promotes plant immunity in Arabidopsis. J. Exp. Bot. 71, 6684–6696 (2020).

39. Z. Wang, et al., Isoprenylation modification is required for HIPP1-mediated powdery mildew resistance in wheat. Plant Cell Environ. 46, 288–305 (2022).

40. H. J. Brabham, et al., Barley MLA3 recognizes the host-specificity effector Pwl2 from *Magnaporthe oryzae*. Plant Cell (2023). 10.1093/PLCELL/KOAD266.

41. D. Gómez De La Cruz, et al., Molecular mimicry of a pathogen virulence target by a plant immune receptor. Science (1979). 392, 1050–1055 (2026).

42. J.-F. Dallery, et al., Gapless genome assembly of *Colletotrichum higginsianum* reveals chromosome structure and association of transposable elements with secondary metabolite gene clusters. BMC Genomics 18, 667 (2017).

43. E. E. Tee, et al., Plasmodesmal closure elicits stress responses. EMBO Rep. (2026). 10.1038/s44319-026-00789-2.

44. K. Aung, et al., Pathogenic Bacteria Target Plant Plasmodesmata to Colonize and Invade Surrounding Tissues. Plant Cell 32, 595–611 (2020).

45. L. Cao, M. C. Blekemolen, N. Tintor, B. J. C. Cornelissen, F. L. W. Takken, The *Fusarium oxysporum* Avr2-Six5 Effector Pair Alters Plasmodesmatal Exclusion Selectivity to Facilitate Cell-to-Cell Movement of Avr2. Mol. Plant 11, 691–705 (2018).

46. M. C. Blekemolen, et al., The primary function of Six5 of *Fusarium oxysporum* is to facilitate Avr2 activity by together manipulating the size exclusion limit of plasmodesmata. Front. Plant Sci. 13 (2022).

47. M. Alazem, S. N. Nuzzi, T. M. Burch-Smith, Regulation of cell-to-cell trafficking by viral movement proteins. J. Exp. Bot. eraf184 (2025). 10.1093/jxb/eraf184.

48. M. Taliansky, L. Torrance, N. O. Kalinina, “Role of Plant Virus Movement Proteins” in Plant Virology Protocols: From Viral Sequence to Protein Function, G. D. Foster, I. E. Johansen, Y. Hong, P. D. Nagy, Eds. (Humana Press, 2008), pp. 33–54.

49. E. E. Tee, S. Samwald, C. Faulkner, “Quantification of Cell-to-Cell Connectivity Using Particle Bombardment” in Plasmodesmata: Methods and Protocols, Y. Benitez-Alfonso, M. Heinlein, Eds. (Springer US, 2022), pp. 263–272.

50. J. Lobstein, et al., SHuffle, a novel *Escherichia coli* protein expression strain capable of correctly folding disulfide bonded proteins in its cytoplasm. Microb. Cell Fact. 11, 753 (2012).

