## Supplementary Figures and Tables for "Immune surveillance at plasmodesmata restricts the movement of a fungal effector"

**This PDF file includes:**

Extended Methods

Figures S1 to S16

Tables S1 to S4

Legends for Datasets S1 to S12

SI References

**Other supporting materials for this manuscript include the following:**

Datasets S1 to S12

### Extended Methods

#### Plant growth

*N. benthamiana* plants for infiltration, or Arabidopsis plants for seed collection or transformation, were grown in a controlled environment room under long-day conditions: 22 °C and 80% relative humidity, with a 16 h light/8 h dark photoperiod. For leaf assays, Arabidopsis plants were grown under short day conditions (22 °C with a 10 h light/14 h dark) in Panasonic Versatile Environmental Test Chambers (MLR-352-PE), equipped with Six Newlec LED T8 colour temperature 4000K and Nine Newlec LED T8 colour temperature 6500K lights. For cultivation of seedlings on sterile media, dry Arabidopsis seeds were sterilised with chlorine gas, placed on growth media (GM) agar plates [4.4 $gL^{-1}$ Murashige Skoog Medium (Duchefa M0222), 0.50 $gL^{-1}$ MES hydrate pH6 (Sigma M8250), 15 $gL^{-1}$ agar (Formedium AGA03)], stratified for 2 days at 5 °C in the dark, then transferred to long-day growth conditions at 20 °C. Rice plants (*Oryza sativa* cv. CO39) were cultivated in 85% humidity, under cycles of 16 h light at 26 °C, and 8 h dark at 24 °C.

#### Construction of DNA plasmids

The coding sequences of *ChEC108*, *HIPP6* and genes encoding other candidate ChEC108-associated proteins were synthesised by Genewiz (Azenta Life Sciences) and cloned into acceptor plasmids with epitope tags and regulatory elements using the Golden Gate system (1). For recombinant protein production in *E. coli*, coding sequences were cloned into pPGN-C or pPGC-K, Golden Gate-compatible backbones derived from pOPIN-F and pOPIN-A, respectively (2). To assemble plasmids for expression in *Magnaporthe oryzae*, the *ChEC108* coding sequence was PCR-amplified with overhangs to allow integration into linearised vector backbones using In-Fusion® cloning. Backbones used were pScBAR with Basta resistance (3) or a custom backbone developed by Yan et al. (4) with sulfonylurea resistance.

To assemble a plasmid for generating the *Colletotrichum higginsianum ΔChec108* strain, 1000 bp genomic regions flanking the *ChEC108* coding sequence were synthesised (Genewiz), placed upstream and downstream of a hygromycin resistance cassette, and cloned into a custom Golden Gate transformation vector named pBin-Δhph-GG derived from pBin-GFP-hph (5). For subsequent transformation of the *ΔChec108* strain, $ChEC108_{pro}:ChEC108:ChEC108_{ter}$ and $ChEC108_{pro}:ChEC108^{D164R}:ChEC108_{ter}$ expression cassettes were assembled by Golden Gate cloning and inserted into pBin-Δhph-GG alongside a $gpdA_{pro}:BAR:trpC_{ter}$ module for Basta selection.

To generate the *hipp6-3* mutant using CRISPR-Cas9 genome editing, 20 bp Cas9 target sites at the *HIPP6* locus were identified using CHOPCHOP (6). Two target sites were selected both upstream and downstream of the coding sequence. sgRNAs were amplified using the template plasmid BCJJ434A (7) with a *U6-26*$_{ter}$ and Golden Gate-compatible overhangs. These were incorporated into Level 1 plasmids with a *U6-26*$_{pro}$, and subsequently into binary Level 2 plasmids alongside *YAO*$_{pro}$*:Cas9-3(intron):E9*$_{ter}$ cassettes, encoded by BCJJ345B (7) and a FastRed selection cassette (8).

#### Agroinfiltration of *Nicotiana benthamiana*

Plasmids were transformed into *Agrobacterium tumefaciens* GV3101 by electroporation, and cultures were diluted in infiltration buffer (in $H_2O$: 10 mM MES pH 5.7, 10 mM $MgCl_2$, 100 µM acetosyringone) prior to infiltration of mature leaves on 3-4-week-old *N. benthamiana* plants. To achieve widespread transformation events across a leaf, bacteria were diluted to an $OD_{600}$ of 0.1-0.4. To achieve discrete single cell transformation events for quantification of protein mobility, bacteria were diluted to an $OD_{600}$ of 0.0001. All infiltration mixtures were prepared with bacteria carrying the RNA silencing suppressor p19 at $OD_{600}$ = 0.1. Leaf material was harvested 2-3 days post infiltration.

#### Bombardment of Arabidopsis

Microprojectile bombardment of fully expanded Arabidopsis leaves, detached from 5-6-week-old rosettes, was carried out as described by Tee et al. (9) using a Biolistic PDS-1000/He particle delivery system (BIO-RAD). Briefly, plasmid DNA (5 µg per construct) was precipitated onto gold microcarriers using $CaCl_2$. Microcarriers were washed and resuspended in 100% ethanol, then applied to carrier discs within microcarrier holders (BIO-RAD), before being fired onto the leaf abaxial surface. Pressure for particle delivery was generated using 1100 psi rupture discs. Bombarded leaves were enclosed within 0.6% or 0.8% agar plates and left in daylight at room temperature for 18-24 h before transformed cells were imaged using confocal microscopy.

#### Transformation of *Magnaporthe oryzae* and inoculation of rice

Transgenic *M. oryzae* strains were generated in the rice pathogenic Guy11 background (10) by protoplast-mediated transformation as described by Talbot et al. (11). Basta- or sulfonylurea-resistant transformants were sub-cultured on Complete Medium (CM) agar (11) at 24 °C under 12 h light/12 h dark photoperiod for 6-8 days. Colonies were flooded with sterile distilled $H_2O$ and gently scraped with an L-shaped cell spreader (E1412-1005, Starlab) to harvest conidia. Conidial suspensions were filtered through one layer of Miracloth (475855, Millipore) then centrifuged at 1200 ×g for 5 min at room temperature. Pelleted conidia were resuspended in 1-2 mL $H_2O$ and diluted to < $1.25 \times 10^5$ conidia/mL using a haemocytometer (Z359629, Merck). Leaf sheaths were peeled from 3-5-week-old rice seedlings and the internal cavity was filled with conidial suspension (12). Inoculated leaf sheaths were placed within petri dishes lined with damp tissue paper and incubated in the dark at 28 °C for 26-27 h prior to confocal imaging.

**Confocal microscopy**

Protein subcellular localisation and cell-to-cell movement were analysed using an LSM Zeiss 800 confocal microscope with Zen software (Blue edition v2.6, Zeiss). Leaves were mounted using double-sided adhesive tape and studied under a 63× (W Plan-Apochromat 63×/1.0 M27, Zeiss) or 20× (W N-Achroplan 20×/0.5 M27, Zeiss) water-dipping objective. To visualise callose deposits at plasmodesmata, immediately prior to imaging, leaves were infiltrated with 0.01% aniline blue (in PBS with 1/10,000 (v/v) Silwet L-77), then rinsed and mounted in $H_2O$ under a VWR® cover slip (631-0120, avantor) and imaged using a 63× (C-Apochromat 63×/1.2 W Corr M27, Zeiss) water-immersion objective. mCherry/mScarlet/tdTomato was excited with a 561 nm diode pumped solid state (DPSS) laser and emission was collected at 560-617 nm. GFP/eGFP/mEGFP was excited with a 488 nm argon laser and emission was collected at 410-546 nm. Aniline blue was excited with a 405 nm UV laser and emission was collected at 410-470 nm.

To quantify protein mobility, bombarded Arabidopsis leaves or agroinfiltrated *N. benthamiana* leaf discs were scanned using a 20× (W N-Achroplan 20×/0.5 M27, Zeiss) water-dipping objective. Due to low signal intensity, movement of ChEC108 fusion proteins in Arabidopsis was studied using a 63× (W Plan-Apochromat 63×/1.0 M27, Zeiss) water-dipping objective. Z-stacks were acquired to cover visible fluorescence at each transformation site, and maximum projections generated in Fiji (13). Nuclear or cell counts were performed manually in Fiji using the Cell Counter plugin. Images were assigned random numbers to hide treatment information from the experimenter during analysis.

To study protein localisation during *M. oryzae* infection of rice, inoculated leaf sheaths were trimmed and mounted epidermis-side-up in water. To visualise nuclei in rice epidermal cells, leaf sheaths were first vacuum infiltrated in DAPI staining solution (10 µg/mL, dissolved in PBS with 0.1% Triton100) and left submerged in staining solution for 30 min before mounting in PBS. Samples were imaged under a VWR® cover slip (631-0120, avantor) using a 63× (C-Apochromat 63×/1.2 W Corr M27, Zeiss) water-immersion objective. DAPI was excited with a 405 nm UV laser and emission was collected at 410-547 nm.

***C. higginsianum* strains and transformation**

The Δ*Chec108* strain was generated in the *ΔChku80* background (14) via targeted homologous recombination to replace the coding sequence of *ChEC108* with a hygromycin resistance cassette, achieved by agrobacterium-mediated transformation of *C. higginsianum* conidia. Complemented lines were subsequently generated in the Δ*Chec108* background by non-targeted insertion of a *ChEC108*$_{pro}$*:ChEC108:ChEC108*$_{ter}$ or *ChEC108*$_{pro}$*:ChEC108*$^{D164R}$*:ChEC108*$_{ter}$ expression cassette, accompanied by a Basta resistance cassette. Constructs were transformed into *A. tumefaciens* AGL1 by electroporation. Cultures were diluted in glycerol induction (GI) broth (in $H_2O$: 1 $gL^{-1}$ $NH_4Cl$, 0.3 $gL^{-1}$ $MgSO_4 \cdot 7H_2O$, 0.15 $gL^{-1}$ KCl, 0.01 $gL^{-1}$ $CaCl_2 \cdot 2H_2O$, 0.0025 $gL^{-1}$ $FeSO_4 \cdot 7H_2O$, 1.8 $gL^{-1}$ glucose, 5 $gL^{-1}$ glycerol, 9.75 $gL^{-1}$ MES/NaOH, 200 µM acetosyringone) to a final $OD_{600}$ of 0.4.

Fungal colonies were cultured at room temperature for 2-3 weeks in the dark on solid Mathur's Medium [in $H_2O$: 2.8 g $L^{-1}$ oxoid mycological peptone, 3.6 g $L^{-1}$ glucose, 1.2 g $L^{-1}$ $MgSO_4$, 2.7 g $L^{-1}$ $KH_2PO_4$, 20 g $L^{-1}$ agar (Formedium AGA03)]. Conidial suspensions (> $2x10^6$ conidia/mL) were collected in GI broth and transformed by mixing with *A. tumefaciens* suspensions in a 1:1 ratio (v/v). Conidial aliquots were spread onto sterile Whatman™ mixed cellulose ester membrane filter discs (47 mm diameter, 7141-114, Cytiva), atop GI agar [GI broth with added 0.69 $gL^{-1}$ $NaH_2PO_4 \cdot H_2O$ and 30 $gL^{-1}$ agar (Formedium AGA03)]. Conidia and bacteria were co-cultivated on GI agar plates at room temperature in the dark for 2 days, then the filter discs were inverted and placed on potato dextrose agar (PDA), supplemented with hygromycin (100 µg/mL) or phosphinothricin (400 µg/mL). After 2 days of dark incubation, filter discs were removed and incubation continued for 4-14 days, or until fungal colonies emerged. Colonies were sub-cultured onto fresh PDA selection plates and incubated for a further 2 weeks. Conidia were harvested as above and 10-fold dilutions spread onto PDA selection plates to obtain colonies derived from single conidia. After 3 days, colonies were sub-cultured on fresh PDA selection plates and conidia harvested after 2-3 weeks, from which a glycerol stock was prepared for long-term storage at –70 °C

##### *C. higginsianum* genotyping

100 mL of potato dextrose broth (PDB) was inoculated with two mycelial plugs, removed from the leading edge of a colony cultured on solid Mathur's Medium as described above. Mycelia were cultured at 27 °C, shaking at 100 rpm for 7 days, then collected by filtration of the media through two layers of Miracloth (475855, Millipore), dabbed dry and frozen in liquid nitrogen. Mycelia were homogenised using a pestle and mortar and gDNA was extracted from 100 mg powder using a DNeasy Plant Mini Kit (QIAGEN), according to the manufacturer's instructions. Transgenes of interest were identified by PCR, using primers listed in *SI Appendix*, Table S4.

##### *C. higginsianum* infection of Arabidopsis

Fungal colonies were cultured on solid Mathur's Medium as described above. Conidial suspensions were collected in water and filtered through a layer of cotton wool within a 5 mL syringe, diluted to $2x10^6$ conidia/mL (for macroscopic lesion measurements) or $2x10^5$ conidia/mL (for microscopic study of infection structures), and used to drop-inoculate the detached leaves of 5-6-week-old Arabidopsis plants, arranged in petri dishes atop 1.5% agar. For macroscopic lesion measurements, avoiding the midrib, one 3 µl droplet of conidial suspension was applied to each leaf. Plates were sealed with at least two layers of parafilm and incubated at 22 °C with a 10 h light/14 h dark photoperiod, covered with one layer of tissue paper to reduce light exposure. Lesions were photographed 6-8 days post-inoculation, and 2D necrotic lesion area measured manually using Fiji. Lesions were excluded from analysis if smaller than 5 $mm^2$ (roughly the size of the applied droplet).

For microscopic study of infection structures, four 1 µL droplets were applied per genotype per leaf, and 6 mm leaf discs were harvested 4 and 5 days post-inoculation. To stain infection structures for brightfield microscopy, leaf discs were boiled with trypan blue staining solution [in $H_2O$: 25% (v/v) glycerol, 25% (v/v) lactic acid, 25% (v/v) water-saturated phenol, 0.025% (w/v) trypan blue] at 95 °C for 2.5 min, while shaking at 500 rpm. Samples were submerged in staining solution for 30 min at room temperature, then rinsed with $H_2O$ and destained in a 5:2 ratio (w/v) chloral hydrate:water with gentle agitation. The chloral hydrate solution was replaced two times throughout the duration of the destaining step (> 3 days). Leaf discs were mounted in 40% glycerol under a VWR® cover slip (631-0120, avantor) and imaged using a Zeiss Axio Imager Z2 with Zen software (Blue edition v2.6, Zeiss), equipped with a 20× (EC Plan-Neofluar 20×/0.5, Zeiss) air-immersion objective and Axiocam 712 colour camera (Zeiss). White balance was adjusted pre-acquisition using the 'pick' function to select a white point for each leaf disc. Brightfield images were acquired as z-stacks with 2 µm intervals between slices, and minimum projections were generated in Fiji. Appressoria and hyphal counts were conducted manually in Fiji using the Cell Counter plugin. Appressoria on trichomes were excluded. Images were assigned random numbers to hide treatment information from the experimenter during analysis.

**Co-immunoprecipitation**

Six leaf discs (8 mm diameter) were harvested from a minimum of three *N. benthamiana* leaves and frozen in liquid nitrogen. Tissue was homogenised using a 2010 Geno/Grinder (SPEX SamplePrep) at 1200 rpm for 1 min, then mixed with 500 µl of IP buffer [in $H_2O$: 50 mM Tris-HCl pH 7.5, 150 mM NaCl, 1 mM EDTA pH 8.0, 10% glycerol, 0.5% NP40 IGEPAL® CA-630, cOmplete™ ULTRA EDTA-free protease inhibitor tablets (1 per 50 mL) (Roche), PhosSTOP EASYpack phosphatase inhibitor cocktail tablets (1 tablet per 50 mL) (Roche), 1 mM $Na_2MoO_4 \times 2H_2O$, 1mM NaF, 5 mM DTT, 1mM PMSF] and rotated end-over-end at 4 °C for 30 min. Extracts were twice centrifuged at 21,130 ×g for 10 min at 4 °C and the supernatant retained. An aliquot of extract was set aside as the input fraction, and the remainder carried forward for IP. To enrich for GFP- or Myc-tagged proteins, 20 µl of GFP-Trap® magnetic agarose beads (Chromotek gtma-20) or 80-100 µl of Pierce™ anti-c-Myc agarose bead slurry (Thermo Scientific 20168) were added to each sample after pre-washing the beads four times in IP buffer. Extracts were rotated end-over-end with beads for 2-5 h at 5 °C. After, beads were washed five times (each for 5 min at 5 °C) in IP buffer. Input and IP samples were boiled with 8-10 µl of loading buffer [100 mM Tris, pH 6.8, 42% (v/v) glycerol, 2.5% (w/v) SDS, 0.04% (w/v) bromophenol blue, 10% (v/v) β-mercaptoethanol] at 95 °C for 15 min, then proteins were fractioned via SDS-PAGE.

**Mass spectrometry**

Immunoprecipitated proteins were separated by SDS-PAGE, and gel slices prepared according to standard procedures described by Schevchenko et al. (15). Peptide aliquots were analysed by nanoLC-MS/MS on an Orbitrap Fusion™ Tribrid™ mass spectrometer coupled to an UltiMate® 3000 RSLCnano LC system (Thermo Scientific). Samples were loaded and trapped using a pre-column, then separated using an analytical column (nanoEase M/Z column, HSS C18 T3, 100 Å, 1.8 µm; Waters). Recalibrated peaklists were generated with MaxQuant 1.6.17.0 (16) using the protein sequence database derived from the *N. benthamiana* draft genome 101, downloaded on 08.02.2019 with 57,140 entries (Sol Genomics Network) plus the Maxquant contaminants database (250 entries) and the custom sequence for ChEC108-GFP. Label-free quantification results from Maxquant were analysed together with search results from an in-house Mascot Server 2.7 (Matrixscience) using the following settings: precursor tolerance of 6 ppm, fragment tolerance of 0.6 Da, enzyme of trypsin/P with a maximum of two allowed missed cleavages, variable modifications: oxidation (M) and acetylation (protein N-terminal), fixed modification: carbamidomethylation (C). Mascot search results were imported into Scaffold 4 (Proteome Software) using identification probabilities of 99% for proteins and 95% for peptides. Data were analysed using SAINTexpress (17) to identify the most likely ChEC108-associated proteins.

**Western blotting**

Proteins within SDS-PAGE gels were transferred to activated polyvinylidene difluoride (PVDF) membranes using the Mini Trans-Blot® Cell transfer system (BIO-RAD). Membranes were incubated for 1-2 h in blocking buffer [5% (w/v) milk, dissolved in Tris-buffered saline with Tween-20 (TBST)] at room temperature, before antibodies (diluted in blocking buffer) were applied: α-GFP (Roche 11814460001, 1/1000 dilution, incubation overnight at 5 °C), α-Mouse-HRP (for use with α-GFP, Sigma-Aldrich A0168, 1/10000 dilution, 6 h incubation at room temperature), α-RFP (abcam ab34771, 1/2000 dilution, incubation overnight at 5 °C), α-Rabbit-HRP (for use with α-RFP, Sigma-Aldrich A0545, 1/20000 dilution, 6 h incubation at room temperature), α-FLAG-HRP (abcam ab49763, 1/5000 dilution, incubation overnight at 5 °C), or α-Myc-HRP (abcam ab62928, 1/5000 dilution, incubation overnight at 5 °C). Western blots were developed using a SuperSignal™ West Femto Trial Kit (Thermo Scientific) and an Amersham™ ImageQuant 500 or 800 Imager. Total protein loading was observed following a 10 s incubation in Coomassie blue staining solution [in $H_2O$: 3.8 mM Coomassie blue,

10% (v/v) acetic acid, 40% (v/v) methanol] and 5-15 min wash in destaining solution [in $H_2O$: 5.2% (v/v) acetic acid, 42% (v/v) methanol].

**Generation of transgenic and mutant Arabidopsis lines**

Plasmids were transformed into GV3101 *A. tumefaciens* by electroporation, and the resulting strains cultured in 10 mL LB aliquots at 28 °C for 16-24 h. 200 mL of LB was inoculated with 2 mL of this starter culture and incubated at 28 ºC for 20-24 h. Bacteria were collected by centrifugation at 5800 ×g for 15 min at room temperature, then resuspended in 200 mL of dipping buffer [0.5× Murashige Skoog Medium, including vitamins (Duchefa Biochemie), 5% sucrose (w/v), 0.0025% (v/v) Silwet L-77]. Arabidopsis inflorescences were dipped in bacterial suspensions for 1 min with gentle agitation, then covered with a dark plastic bag for 24 h.

To generate Arabidopsis lines constitutively expressing fluorescent proteins (*ChEC108-eGFP* or *mEGFP-HIPP6* and their mutant variants), T1 seed were collected and screened for chemical resistance on sterile GM agar plates supplemented with kanamycin (50 µg/mL) or phosphinothricin (10 µg/mL). Surviving individuals were transplanted to soil and screened for fluorescence via confocal microscopy. To generate the *hipp6-3* mutant, fluorescent T1 seeds carrying the FastRed selection marker were identified using a Zeiss Lumar.V12 stereoscope, equipped with a 1.2× objective, 10× eyepiece lens, BP545/25 excitation filter and BP605/70 emission filter. T2 plants were genotyped by PCR from gDNA to identify individuals homozygous for *HIPP6* deletion (*SI Appendix*, Fig. S8). *hipp6-3* plants used for experimentation originated from transgene-free T3 seeds or subsequent generations.

**Protein production from *Escherichia coli***

To produce proteins for structural studies, plasmids were transformed into SHuffle *E. coli* (18) and the resulting strains cultured in 50 mL LB for 16 h at 30 ºC. 8 L LB was inoculated with this starter culture (5 mL/L) and incubated at 30 ºC until the $OD_{600}$ reached 0.6-0.8. Expression was induced at 18 ºC by adding 1 mM isopropyl β-D-1-thiogalactopyranoside (IPTG). Induction was left to proceed for 17-24 h before cells were pelleted by centrifugation at 7500 ×g for 10 min at 4 ºC. Pellets from each 1 L culture were resuspended in 200 mL of immobilised metal affinity chromatography (IMAC) buffer [in $H_2O$: 150 mM Tris-HCl pH 8.0, 500 mM NaCl, 5% (v/v) glycerol, 50 mM glycine, 20 mM imidazole], supplemented with cOmplete™ EDTA-free protease inhibitor cocktail tablets (Sigma-Aldrich). Cells were lysed by sonication (50% amplitude, 1 s on/3 s off, 8 min pulse duration) using a Vibra Cell™ VC505 equipped with a 13 mm probe (SONICS). Lysate was centrifuged at 45,000 ×g for 30 min at 4 ºC, then the supernatant loaded onto a 5 mL HisTrap™ HP NTA column (Cytiva), equilibrated in IMAC buffer, for enrichment of 6×His-tagged fusions. After elution in IMAC buffer supplemented with 500 mM imidazole, proteins were separated by size exclusion chromatography (SEC) using a Superdex™ 75 HiLoad™ 26/600 column (Cytiva) equilibrated in SEC buffer (in $H_2O$: 20 mM HEPES pH 7.5, 150 mM NaCl). Fractions containing desired proteins were identified using SDS-PAGE, pooled and incubated with 6×His-tagged human rhinovirus 3C protease (10 µg/mg of desired protein) for 12-16 h. 3C protease, cleaved affinity tags and remaining uncleaved contaminants were removed by IMAC using 5 mL HisTrap™ columns equilibrated in SEC buffer. Fractions containing desired proteins were identified using SDS-PAGE, pooled, then concentrated to 5 mL using VivaSpin® 20 mL concentrators (Sartorius) with a 3 kDa, 10 kDa, or 30 kDa molecular weight cut-off, depending on the size of the protein or complex. Samples were further purified by loading onto Superdex™ 75 HiLoad™ 16/600 columns (Cytiva), equilibrated in SEC buffer. Fractions containing desired proteins and minimal contamination were identified using SDS-PAGE, pooled and concentrated as above, then frozen in liquid nitrogen and stored at −70 °C.

For in vitro binding assays, proteins were purified according to Yu et al. (19). Briefly, this includes expression of proteins in SHuffle *E. coli* with an N-terminal 6×His or 6×His-GB1 fusion tag, lysis by sonication followed by IMAC and desalting. N-terminal tags were cleaved with 3C protease for 16 h at 4 ºC. The cleaved protein of interest was separated from the N-terminal fusion tag, any uncleaved protein, and 6×His-tagged 3C protease using IMAC and subsequently purified further by SEC. Proteins were concentrated using a 3 kDa or 10 kDa molecular weight cut-off Amicon centrifugal concentrator (Millipore Sigma), frozen in liquid nitrogen and stored at −70 °C.

#### Protein crystallisation

Initial crystallisation trials were conducted using the sitting drop vapour diffusion method with commercially available sparse matrix screens, dispensed into MRC 96-Well 2-Drop Plates (Molecular Dimensions). Proteins were defrosted on ice and centrifuged at 21,000 ×g for 10 min at 4 ºC to pellet precipitants. 0.3 µL of protein (10 or 20 mg/mL) was mixed with 0.3 µL of precipitant solution and the resulting drop dispensed using an Oryx8 Protein Crystallization Robot (Douglas Instruments) alongside a 40 µL reservoir of concentrated precipitant solution. Plates were immediately sealed with ClearVue adhesive seals (Molecular Dimensions) and stored at 18 ºC. Drops were monitored periodically over 1-2 months using a Rock Imager® (Formulatrix).

The best morphology for ChEC108-HMA1 complex crystals was observed in 25% Polyethylene glycol monomethyl ether (PEG-MME) 2K, 0.1 M HEPES pH 7.5 (well B6, PEGs Suite, QIAGEN) using an initial protein concentration of 20 mg/mL, and further optimisation was not required. Initial screens yielded ChEC108-HMA2 complex crystals under multiple conditions, including 25% PEG 6K, 0.1 M TRIS pH 8.5 (well D9, PEGs Suite, QIAGEN), and optimisation screens were designed around this hit. The best morphology for ChEC108-HMA2 crystals was observed in 16% PEG 6K, 0.1 M TRIS pH 7.5, using an initial protein concentration of 20 mg/mL. Prior to X-ray diffraction, crystals were cryoprotected using 20% ethylene glycol or a solution of 10% glycerol and 10% ethylene glycol (dissolved in precipitant solution from the relevant reservoir), then mounted onto X-ray-transparent loops and vitrified in liquid nitrogen.

**X-ray crystallography and data processing**

X-ray diffraction data were collected at beamline i24 (MX32728-3 for ChEC108-HMA1) and i04 (MX32728-5 for ChEC108-HMA2) of the Diamond Light Source synchrotron facility (Oxford, UK). Data were auto-processed via the xia2 dials pipeline (20), accessed via Diamond ISPyB. Scaled, unmerged data were imported into ccp4i2 v.8.0 (21) and processed using *AIMLESS* (22). Molecular replacement was carried out in *Phaser* (23). Regarding ChEC108-HMA2 complexes, the predicted structures of ChEC108 and HMA2 monomers were obtained using AlphaFold2 (24), accessed via the Google Colaboratory and ColabFold (25), and used as search models for molecular replacement. Regarding ChEC108-HMA1 complexes, the ChEC108 and HMA2 crystal structures solved prior were used as search models. *Bucaneer* (26) was used for automatic model building, then further manual building and refinement were carried out in Coot (27). Cycles of manual and automatic refinement were carried out in *REFMAC5* (28). X-ray data collection and refinement statistics are given in *SI Appendix,* Tables S1 and S2. X-ray fluorescence scanning was conducted using crystalline material containing ChEC108-HMA2 complexes at beamline i04 (MX32728-16) at Diamond Light Source. Crystals were exposed to high energy (13,000 eV) incident X-rays, and the energy of emitted X-rays quantified to indicate the presence of heavy elements.

**Isothermal titration calorimetry**

In vitro purified proteins (ChEC108, ChEC108$^{D164R}$ or HMA domains) were diluted in SEC buffer (in $H_2O$: 20 mM HEPES, pH 7.5, 150 mM NaCl) supplemented with 1 mM DTT and 50 µM $ZnCl_2$, then loaded into the experimental cell (250 µL of HMA domain) or syringe (50 µL of ChEC108 or ChEC108$^{D164R}$) of a MicroCal PEAQ-ITC machine (Malvern, UK). Protein concentrations in the cell/syringe were 20/200 µM for all HMA/ChEC108 combinations except for HIPP6-HMA2/ChEC108, where concentrations of 30/300 µM were used. Each run was conducted at 25 °C and involved a single 0.5 µL injection from the syringe, followed by 18 injections of 2 µL each at 120 s intervals. ITC experiments were performed in triplicate. Raw titration data were uploaded to AFFINIMeter (29), where baselines were corrected and binding parameters calculated.

**Analytical size exclusion chromatography**

In vitro purified proteins (ChEC108, ChEC108$^{D164R}$, HMA1, HMA2) were incubated on ice for at least 1 h, either alone or in pairs. Proteins were separated on a Superdex 75 Increase 5/150 GL (Cytiva) column pre-equilibrated in SEC buffer (in $H_2O$: 20 mM HEPES, pH 7.5, 150 mM NaCl), supplemented with 1 mM DTT and 50 µM $ZnCl_2$. Column eluate was monitored by measuring absorbance at 280 nm.

**RNA extraction from Arabidopsis**

Seven 19-day-old whole seedlings were collected from sterile GM agar plates, equating to 70-90 mg of fresh weight per sample. Three replicate samples, harvested from separate plates, were collected per genotype. Seedlings were frozen in liquid nitrogen and homogenised using a 2010 Geno/Grinder (SPEX SamplePrep) at 1200 rpm for 1 min. RNA was extracted using the RNeasy Plant Mini Kit (QIAGEN) using buffer RLT according to the manufacturer's instructions with the following modification: after tissue homogenisation using the QIAshredder column, two or three centrifugation steps were performed to maximise removal of debris. RNA extracts were DNase-treated using the TURBO DNA-freeTM Kit (Invitrogen) according to the manufacturer's rigorous procedure.

**Transcriptomic analysis**

Total RNA samples were submitted to Genewiz for library preparation with polyA selection and standard RNA-sequencing. Illumina sequencing generated 150 bp paired-end reads and was carried out at a depth of 20 million reads per sample. Adaptors and low-quality reads (Phred score $< 20$) were trimmed using TrimGalore v0.5.0 (30) and Cutadapt v3.4 (31). Pre- and post-trimming, data quality was assessed using FastQC v0.11.8 (32). Reads were mapped to the Arabidopsis reference genome (TAIR10; NCBI Refseq GCA_000001735.1) using HISAT2 v2.1.0 (33). Mapping quality was assessed using and MultiQC v.1.7 (34) before data were imported into Seqmonk v.1.49.0 (Babraham Bioinformatics) for visualisation and quantification. Replicate sets were assigned, and the default RNA-seq quantitation pipeline in Seqmonk was used to generate raw counts. Genes showing significant differential expression between genotypes were identified using Wald tests within the R package, *DEseq2* v1.48.2 (35) using cutoffs of $\log_2$ fold change $\geq |1|$ and adjusted p value $\leq 0.05$ to define differential expression. Genes with less than 10 counts summed across all samples were excluded. GO enrichment analysis was conducted using the R package, *clusterProfiler* v4.16.0 (36), with p and q value cutoffs of $< 0.05$.

**Statistical analysis and data presentation**

All statistical tests were conducted in R. Details to accompany each figure are given in Dataset S12. Bootstrapping analysis of mobility assay data was performed using the *medianBootstrap* function, as described by Johnston & Faulkner (37). Packages *emmeans* v2.0.0 (38), *lme4* v1.1-37 (39) and *lmerTest* v3.1-3 (40) were used to generate linear mixed effect models. All numerical data were plotted using *ggplot2* v4.0.0 (41). Venn diagrams were drawn using *ggVennDiagram* v1.5.4 (42). Predicted and crystal structures of proteins were visualised using ChimeraX v1.9 (43).

### Supplementary Figures

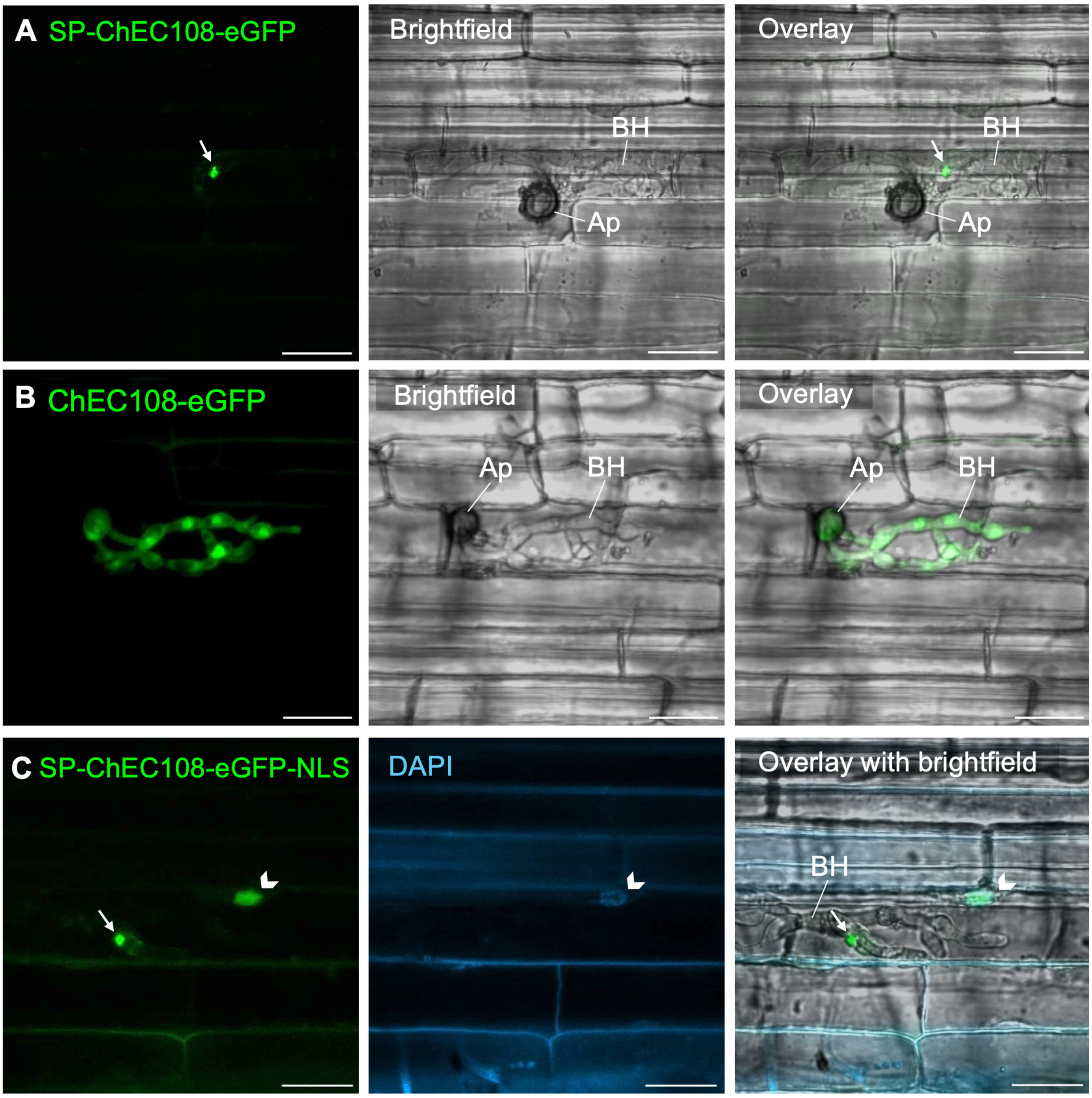


**Figure S1. ChEC108 resembles a cytoplasmic effector when expressed heterologously in *Magnaporthe oryzae*. A-C.** Rice leaf sheath epidermal cells inoculated with *M. oryzae* expressing (A) *AVR-PikD*$_{pro}$*:SP-ChEC108-eGFP* with the native N-terminal signal peptide (SP) included, (B) *AVR-PikD*$_{pro}$*:ChEC108-eGFP* with the SP absent, or (C) *AVR-PikD*$_{pro}$*:SP-ChEC108-eGFP-NLS,* including a non-native triple nuclear localisation signal (NLS) extension. Concentration of eGFP signal at the BIC (arrows) can be seen in A and C only. SP-ChEC108-eGFP-NLS was also observed in host cell nuclei, marked by DAPI staining in C (arrowhead). Ap = appressorium, BH = biotrophic hypha. Images were acquired 26-30 hours post-inoculation and are representative of at least two independent *M. oryzae* strains generated to express each fusion protein. Scale bar = 20 µm.

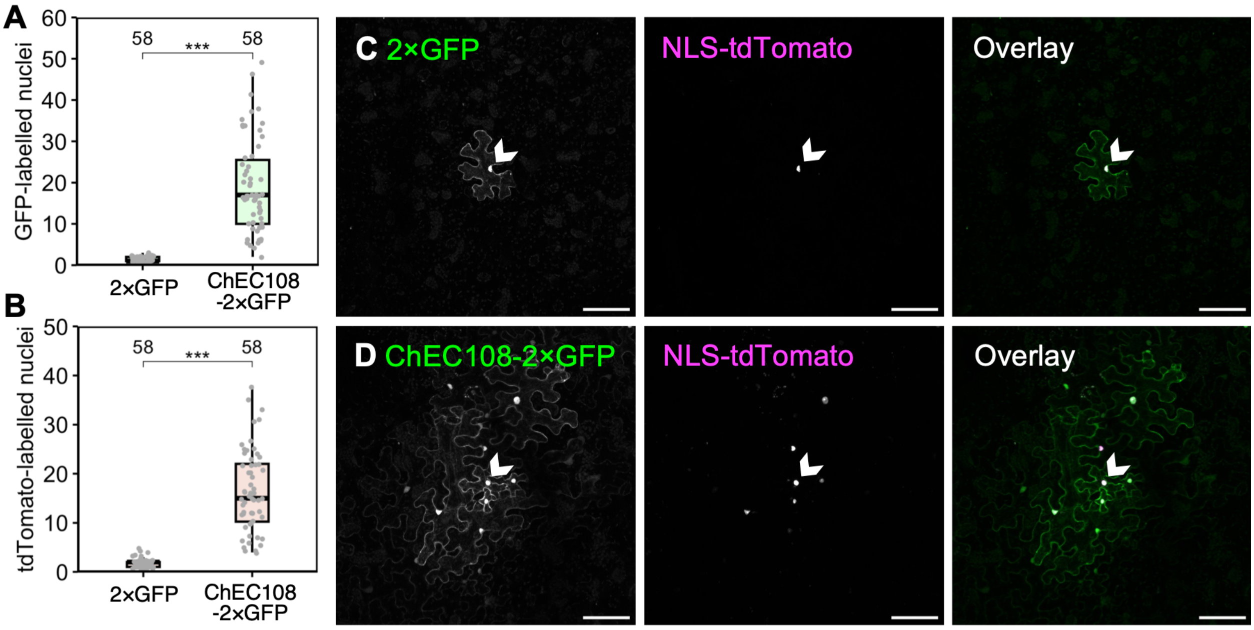


**Figure S2. ChEC108 is mobile and promotes cell-to-cell trafficking in *N. benthamiana.*** *$35S_{pro}$:NLS-tdTomato* transiently expressed in single *N. benthamiana* leaf epidermal cells via low-OD agroinfiltration alongside *$35S_{pro}$:ChEC108-2×GFP* or *$35S_{pro}$:2×GFP* as a control. **A-B.** Mobility of GFP (A) or tdTomato signal (B) quantified 3 days post-infiltration by counting the number of fluorescent nuclei at transformation sites. Numeric annotations indicate sample size (sites), collected across 3 leaves per treatment, with a maximum of 20 sites per leaf. *** indicates $p < 0.001$, determined by bootstrapping analysis with 5000 iterations. **C-D.** Representative transformation sites, showing 2×GFP confined to a single cell (C), while ChEC108-2×GFP signal can be seen spreading to adjacent cells (D). Spread of NLS-tdTomato signal is enhanced in D vs C. Arrowheads indicate the nuclei of transformed cells. Scale bar = 100 μm.

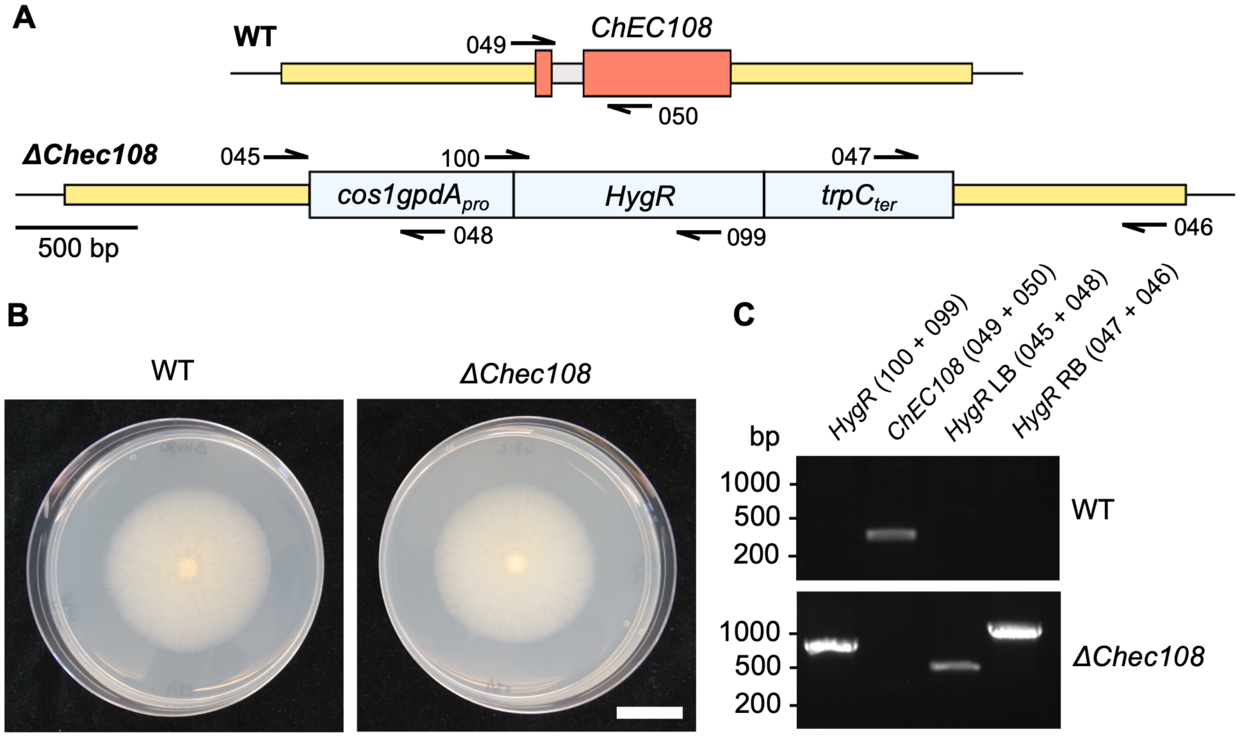


**Figure S3. Generation of *ΔChec108 C. higginsianum* by homologous recombination. A.** Schematic representing wild-type (WT) and *ΔChec108* genomic sequences at the *ChEC108* locus in the *ΔChku80* background. In *ΔChec108*, the *ChEC108* coding sequence is replaced by a hygromycin resistance gene (*HygR*), under control of an *Aspergillus nidulans cos1gpdA*$_{pro}$ and *trpC*$_{ter}$. Homologous recombination to integrate the *HygR* cassette was guided by 5' and 3' flanking regions (yellow). **B.** Comparable growth of WT and *ΔChec108* strains on solid Mathur's medium plates 8 days post-revival from a glycerol stock. Scale bar = 2 cm. **C.** Genotyping of *ΔChec108* by PCR from gDNA. Numbers represent primers used, the binding sites of which are depicted in A (not to scale). LB = left border, RB = right border. Primer sequences are given in *SI Appendix,* Table S4.

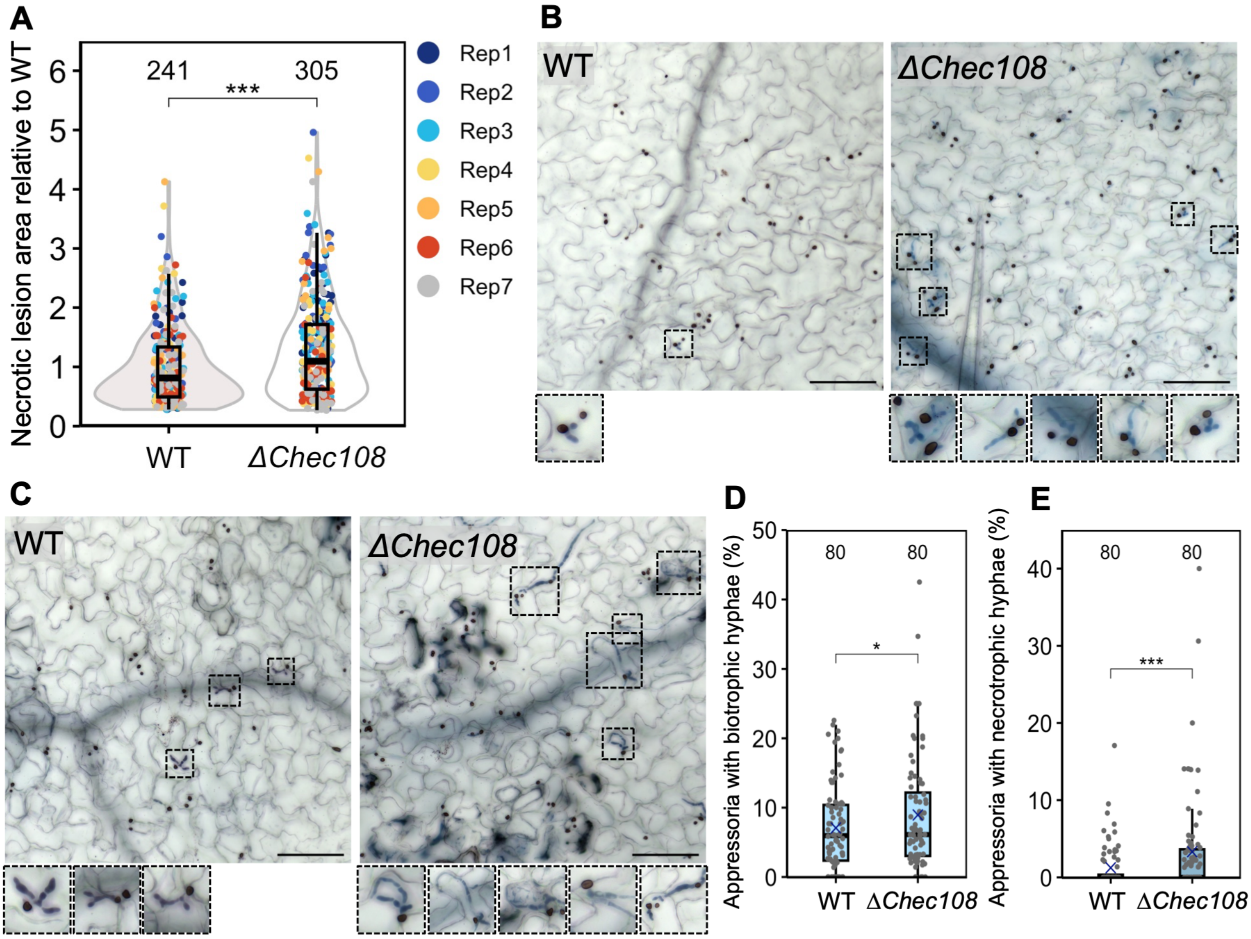


**Fig. S4. Loss of *ChEC108* enhances *C. higginsianum* virulence. A.** Area of necrotic lesions induced by WT or *ΔChec108 C. higginsianum* on Col-0 Arabidopsis leaves. Data were normalised by dividing all measurements by the mean area of WT lesions for a given replicate (Rep). Numeric annotations indicate sample size (lesions). *** indicates $p < 0.001$, determined using a linear mixed model, with replicate as a random effect and fungal genotype as a fixed effect. **B-C.** Col-0 Arabidopsis leaves infected with WT or *ΔChec108 C. higginsianum,* stained with trypan blue 4 days (B) or 5 days (C) post-inoculation. Appressoria and biotrophic hyphae are visible within epidermal cells. Scale bar = 100 µm. Lower panels show a magnified view of hyphae within regions indicated by black boxes. **D-E.** Percentage of appressoria associated with biotrophic hyphae (D) and necrotrophic hyphae (E) 5 days post-inoculation. Numeric annotations indicate sample size (images), collected from 10 leaves, with 8 images captured across 3-4 inoculated zones per genotype per leaf. * indicates $p < 0.05$ and *** indicates $p < 0.001$, determined by a binomial general linear mixed model with genotype as a fixed factor, alongside leaf and zone as nested random factors. The percentage of appressoria associated with biotrophic and necrotrophic hyphae 4 days post-inoculation is shown in Fig. 1.

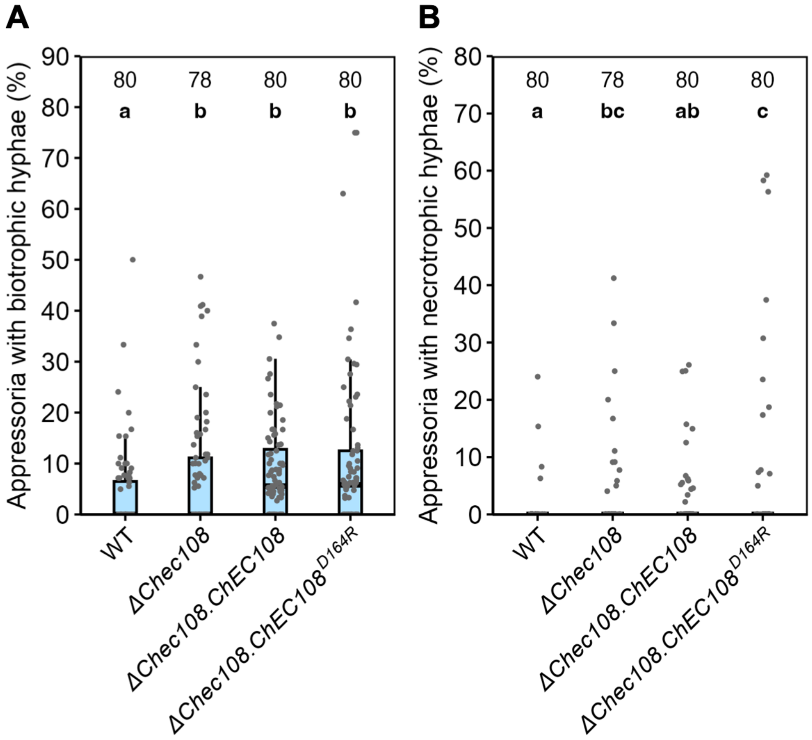


**Figure S5. Enhanced virulence of *ΔChec108* is partially complemented by re-introduction of *ChEC108*, but not *ChEC108*$^{D164R}$. A-B.** Percentage of appressoria of WT, *ΔChec108, ΔChec108.ChEC108 or ΔChec108.ChEC108*$^{D164R}$ *C. higginsianum* associated with trypan blue-stained biotrophic hyphae (A) and necrotrophic hyphae (B) 4 days post-inoculation of Arabidopsis leaves. Numeric annotations indicate sample size (images), collected from 10 leaves, with 8 images captured across 3-4 inoculated zones per genotype per leaf. Compact letters indicate significant differences ($p < 0.05$), determined by a post-hoc Tukey test on a binomial general linear mixed model with genotype as a fixed factor, alongside leaf and zone as nested random factors.

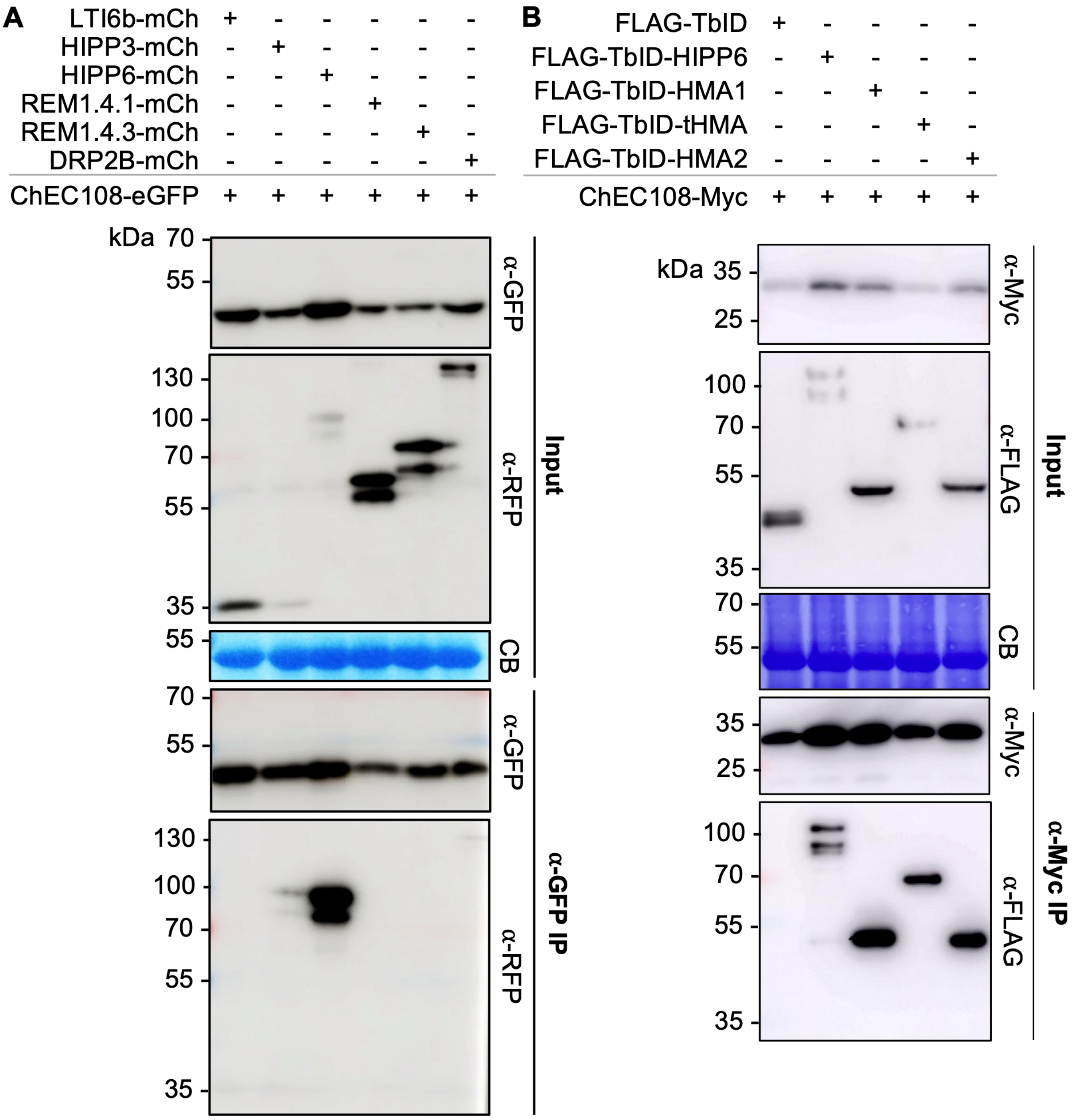


**Figure S6. ChEC108 associates with HIPP6 HMA domains in *N. benthamiana*. A.** Co-IP demonstrating association of ChEC108-eGFP with mCherry (mCh)-tagged HIPP6. LTI6b-mCh served as a negative control. Association with other mCh-tagged candidate interactors was not reproducibly detected. **B.** Co-IP demonstrating association of ChEC108-4×Myc with 3×FLAG-TurboID (FLAG-TbID)-tagged HMA1, HMA2 and tHMA. FLAG-TbID alone and FLAG-TbID-HIPP6 served as negative and positive controls, respectively. CB = Coomassie Blue staining of the Rubisco large subunit. Constructs were transiently expressed in *N. benthamiana* leaves under a $35S_{pro}$. Blots are representative of three independent replicates.

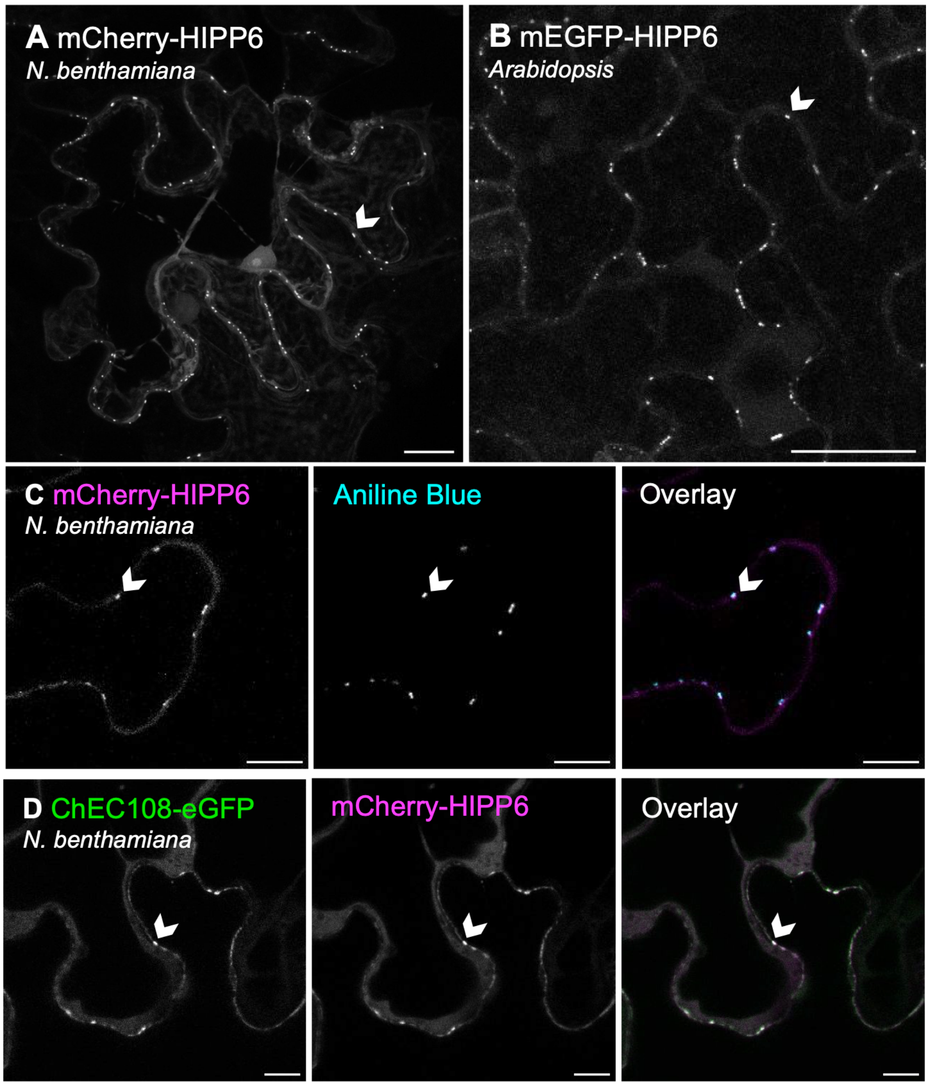


**Figure S7. HIPP6 localises to plasmodesmata in *N. benthamiana* and Arabidopsis. A.** *35S$_{pro}$:mCherry-HIPP6* transiently expressed in *N. benthamiana* leaf epidermal cells via agroinfiltration. Scale bar = 20 µm. **B.** *AtUBQ10$_{pro}$:mEGFP-HIPP6* stably expressed in Arabidopsis leaf epidermal cells. Scale bar = 20 µm. **C-D.** Co-localisation of mCherry-HIPP6 with aniline blue-stained callose deposits (C) or ChEC108-eGFP, introduced by co-agroinfiltration (D). Scale bar = 10 µm. Arrowheads indicate plasmodesmata.

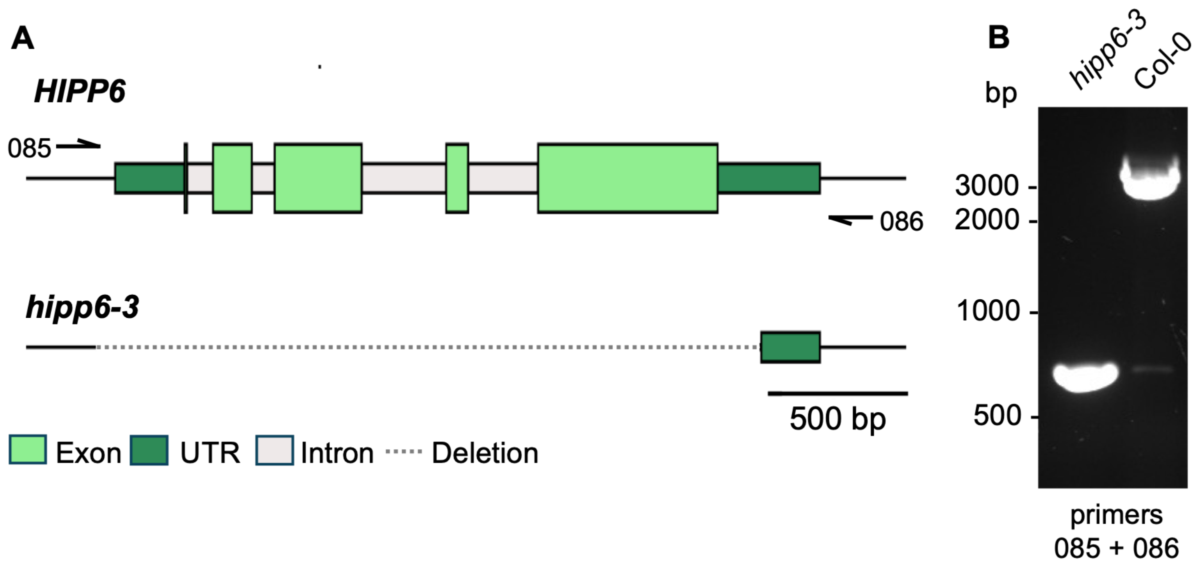


**Figure S8. Generation of *hipp6-3* Arabidopsis using CRISPR-Cas9. A.** Schematic representing wild type *HIPP6* and *hipp6-3* genomic sequences. *hipp6-3* lines were generated by simultaneously targeting Cas9 to the promoter and 3’ UTR, resulting in a 2367 bp deletion encompassing the 5’ UTR and all exons and introns. **B.** Genotyping of *hipp6-3* plants by PCR from gDNA. Numbers represent primers used, the binding sites of which are depicted in A (not to scale). Primer sequences are given in *SI Appendix,* Table S4.

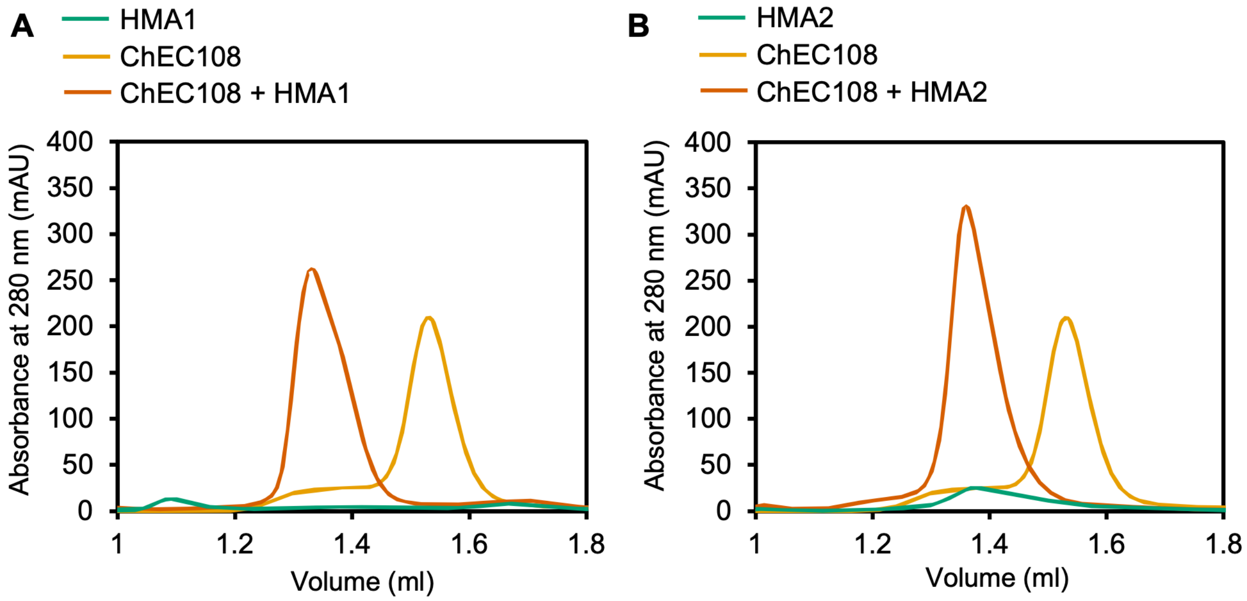


**Figure S9. ChEC108-HMA1 and ChEC108-HMA2 complex formation captured in vitro by aSEC. A-B.** Size exclusion chromatograms showing a leftward peak shift when ChEC108 was mixed with HMA1 (A) or HMA2 (B) in the presence of 1 mM DTT and 50 µM $ZnCl_2$ (orange trace). Yellow and green traces are chromatograms representing ChEC108 and HMA domains alone, respectively.

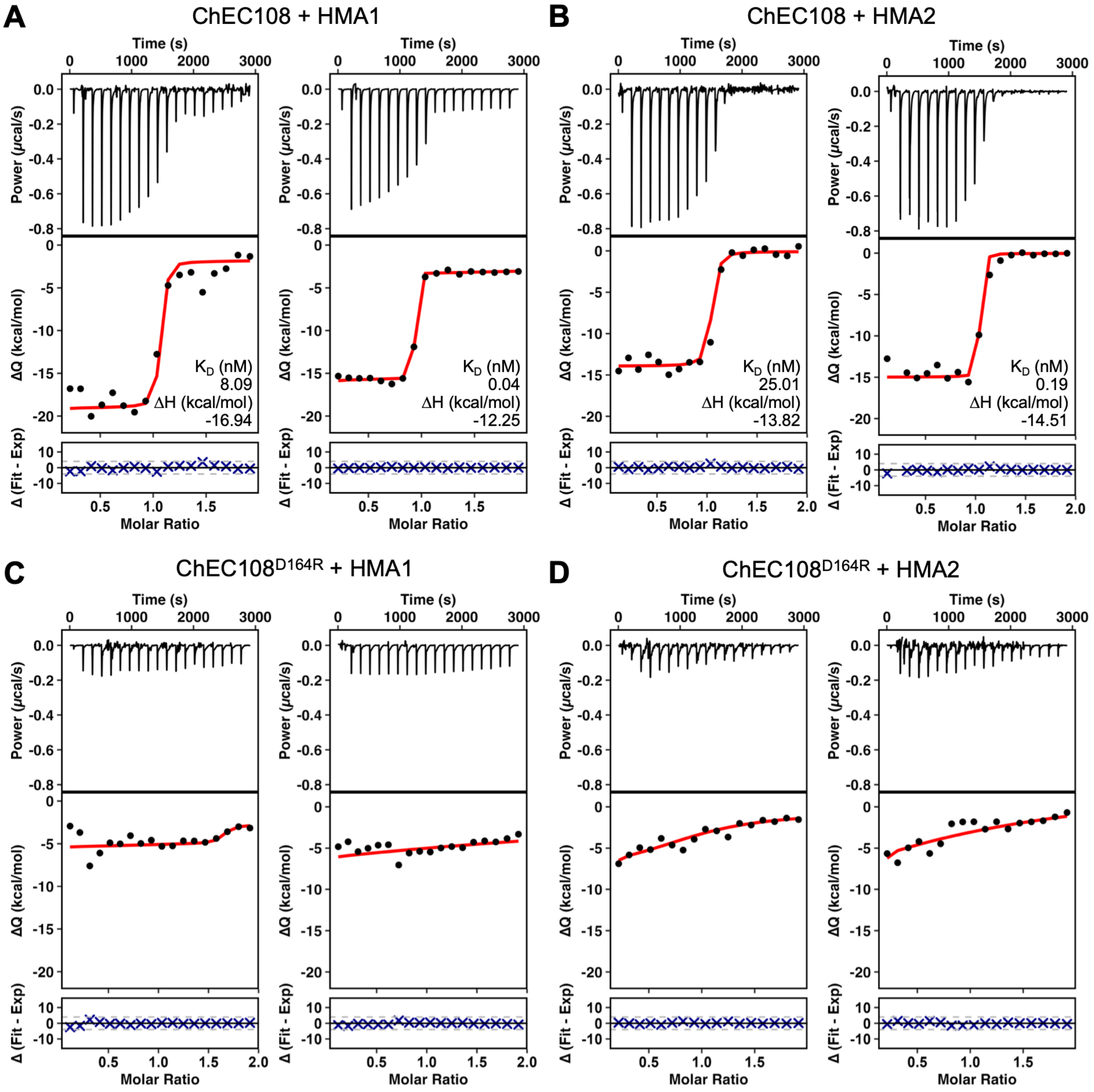


**Figure S10. ITC revealed HMA1 and HMA2 are tightly bound by ChEC108 in vitro, but not by ChEC108$^{D164R}$. A-B.** In vitro characterisation of ChEC108-HMA1 (A) or ChEC108-HMA2 binding (B) by ITC. **C-D.** Binding of ChEC108$^{D164R}$ to HMA1 (C) or HMA2 (D) was not detected. In each panel, left and right traces represent two independent replicates, with an additional third replicate displayed in Fig. 4.

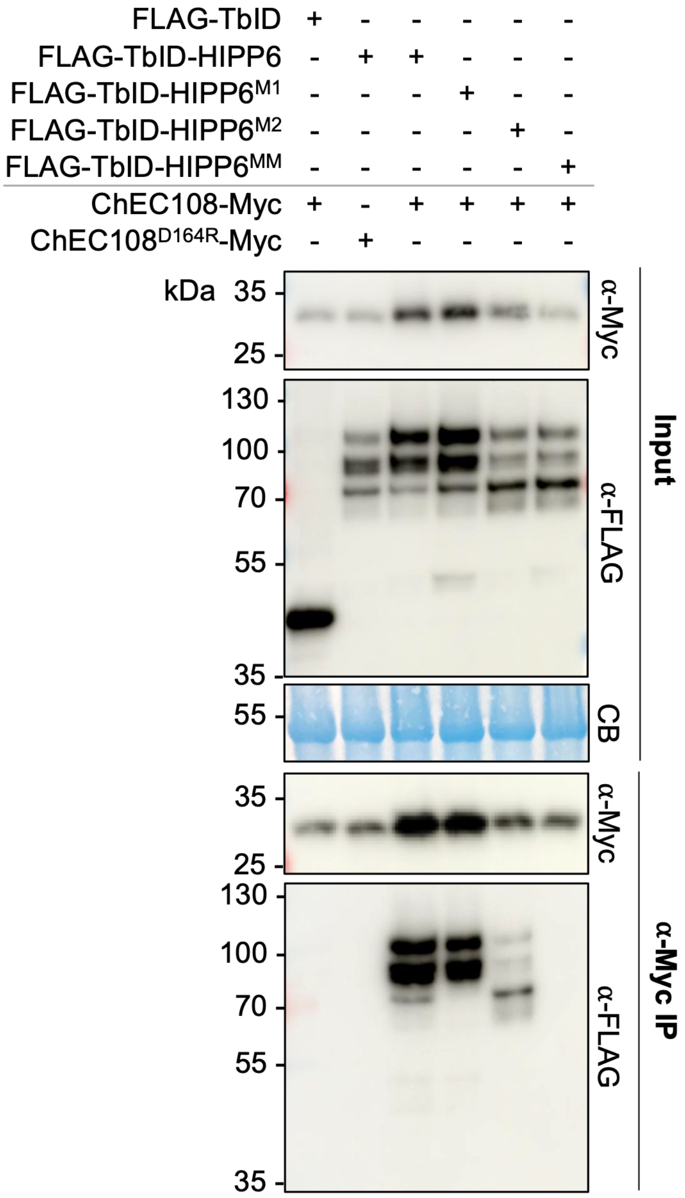


**Figure S11. Asp164 of ChEC108 and the CXXC motifs of HIPP6 are critical for ChEC108-HIPP6 binding in vivo.** Co-IP demonstrating D164R mutation of ChEC108 abolished association with wild-type HIPP6. Although HIPP6$^{M1}$ (carrying a C$^{34}$XXC to G$^{34}$XXG mutation in HMA1) and HIPP6$^{M2}$ (carrying a C$^{164}$XXC to G$^{164}$XXG mutation in HMA2) associated with ChEC108, HIPP6$^{MM}$ (in which both CXXC motifs are mutated to GXXG) did not. Bait proteins were tagged with 4×Myc. Prey proteins were tagged with 3×FLAG-TurboID (FLAG-TbID). FLAG-TbID alone and FLAG-TbID-HIPP6 were included as negative and positive controls, respectively. CB = Coomassie Blue staining of the Rubisco large subunit. Constructs were transiently produced in *N. benthamiana* leaves under a *35S*$_{pro}$. Representative of three independent replicates.

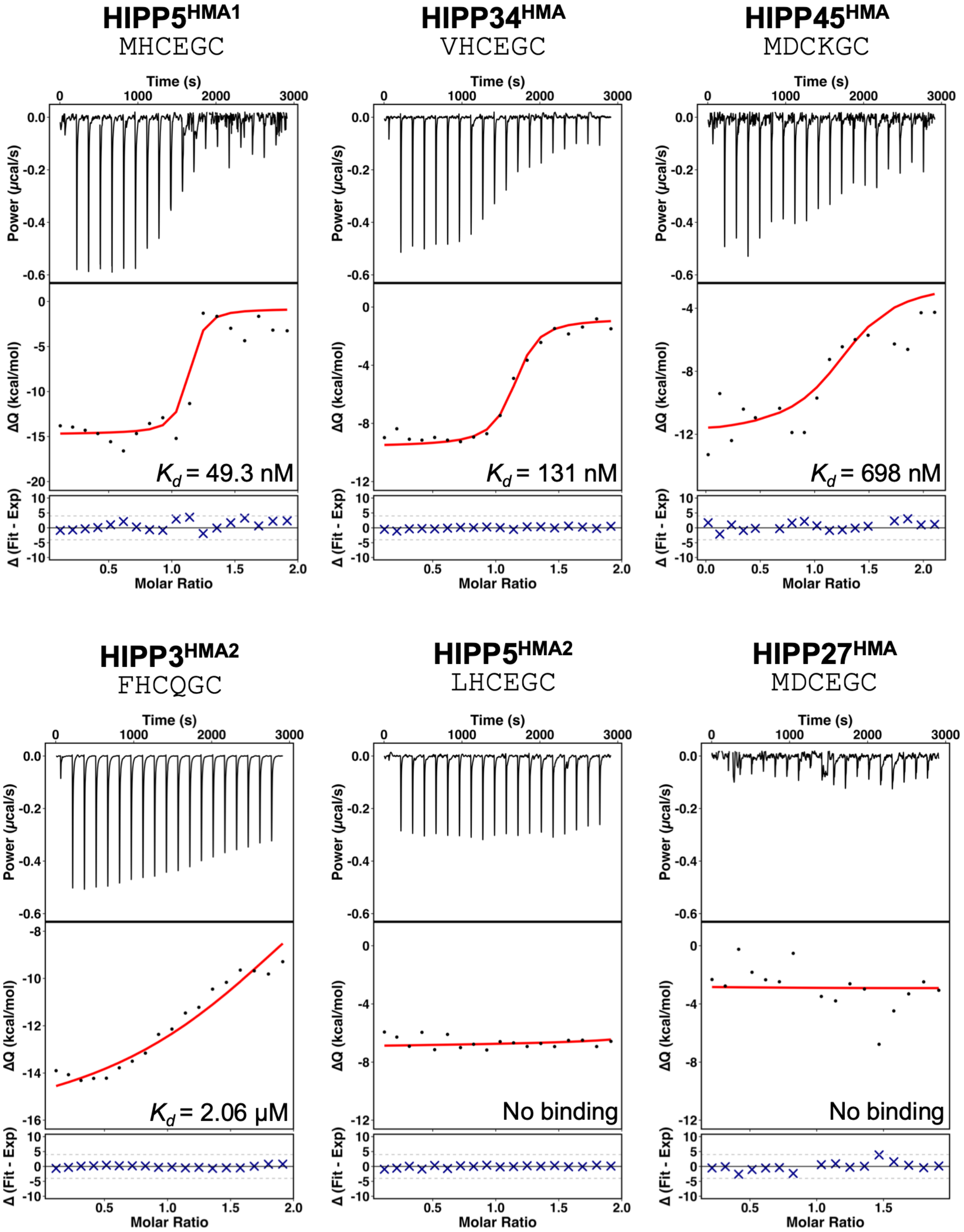


**Figure S12. ITC revealed HMA domains of different HIPPs can bind ChEC108. A.** In vitro characterisation of ChEC108-HMA binding, including HMA1 from HIPP5, HMA2 from HIPP3, and the HMA domains from HIPP34 and HIPP45 by ITC. Binding of ChEC108 to HMA2 from HIPP5 and the HMA domain from HIPP27 was not detected. Metal-binding (M/LXCXXC) motifs of each HMA domain are displayed. For reference, HMA1 and HMA2 from HIPP6 contain motifs reading MHCEGC and LHCEGC, respectively.

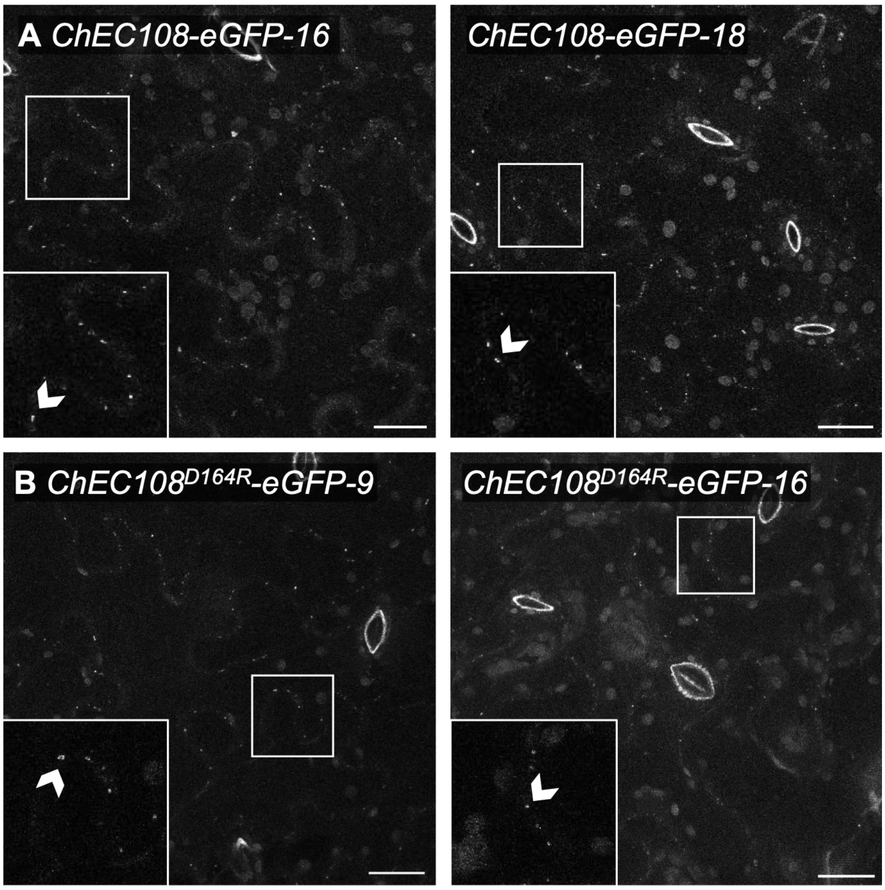


**Figure S13. ChEC108 targets plasmodesmata independent of HIPP6 when stably expressed in Arabidopsis. A-B.** Plasmodesmata-localised eGFP signal in leaf epidermal cells of two independent transgenic Arabidopsis lines expressing *AtUBQ10*$_{pro}$*:ChEC108-eGFP* (A) or *AtUBQ10*$_{pro}$*:ChEC108*$^{D164R}$*-eGFP* (B). Chloroplast- and stomata-associated autofluorescence are also visible. Inset panels show a magnified view of areas indicated by white boxes. Arrowheads indicate plasmodesmata. Scale bar = 20 µm.

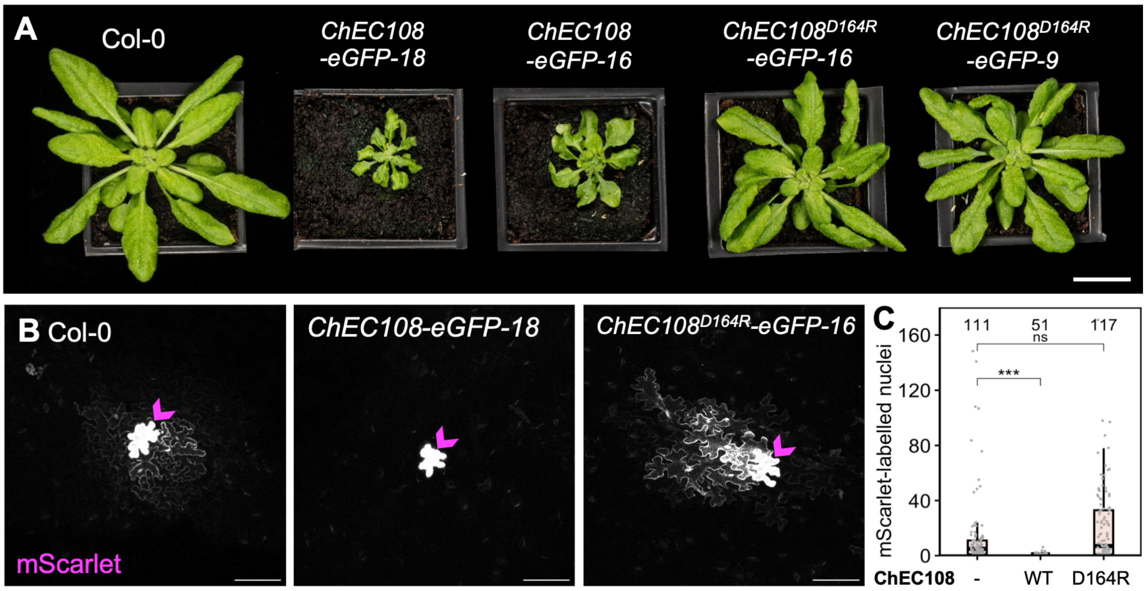


**Figure S14. Cell to cell connectivity is reduced upon ChEC108-HIPP6 binding. A.** 6-week-old rosettes of Col-0 Arabidopsis, and lines constitutively expressing *AtUBQ10*$_{pro}$*:ChEC108-eGFP* or *AtUBQ10*$_{pro}$*:ChEC108*$^{D164R}$*-eGFP*. Scale bar = 2 cm. **B.** mScarlet introduced via bombardment of Col-0, *AtUBQ10*$_{pro}$*:ChEC108-eGFP* or *AtUBQ10*$_{pro}$*:ChEC108*$^{D164R}$*-eGFP* lines. Images are maximum projections covering representative bombardment sites. Magenta arrowheads indicate bombarded cells. Scale bar = 100 µm. **C.** Mobility of mScarlet, quantified by counting the number of fluorescent nuclei at transformation sites. Numeric annotations indicate sample size (sites), collected across 8-10 leaves per genotype, with a maximum of 20 sites per leaf. *** indicates $p < 0.001$ and ns indicates no significant difference, determined by bootstrapping analysis with 5000 iterations.

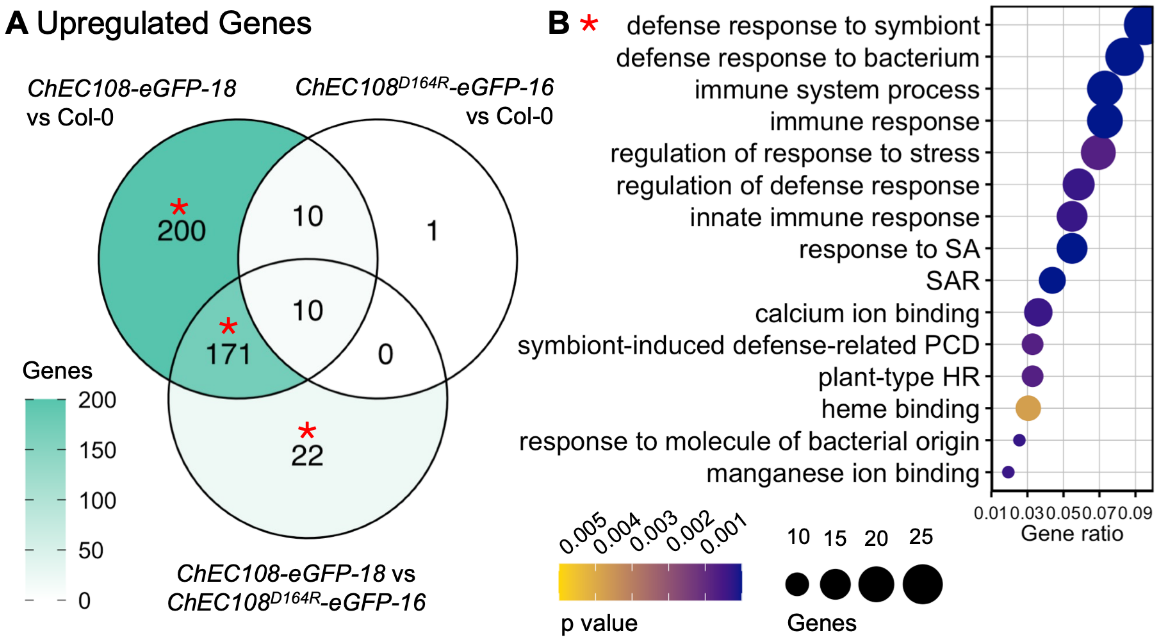


**Figure S15. Constitutive expression of *ChEC108* triggers defence-related transcriptomic responses. A.** Transcriptomic analysis of Arabidopsis seedlings of indicated genotypes. Venn diagram shows overlap of genes differentially expressed (upregulated) between genotypes ($\log_2$ fold change ≥ |1|, Wald test adjusted p value ≤ 0.05). Asterisks indicate sets of genes induced in the presence of ChEC108-eGFP. **B.** Top 15 enriched GO terms assigned to differentially expressed genes in sectors marked by asterisks in A. p values were determined by one-sided Fisher's exact tests and adjusted using the Benjamini-Hochberg method. HR = hypersensitive response, PCD = programmed cell death, SA = salicylic acid, SAR = systemic acquired resistance.

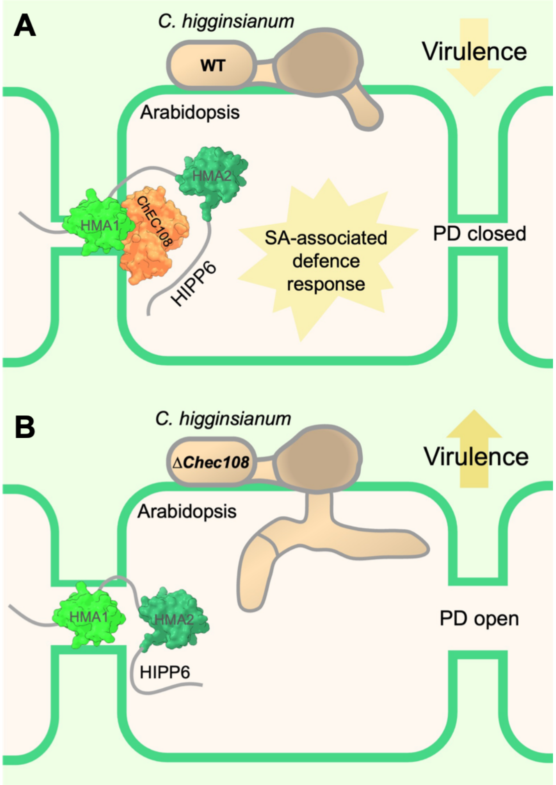


**Figure S16. Model depicting the consequences of ChEC108-HIPP6 interaction. A.** Upon *C. higginsianum* infection of Arabidopsis, ChEC108 is secreted, enters the host cytoplasm and localises to plasmodesmata. Here, ChEC108 binds HIPP6 via direct contact with HMA1 or HMA2. This triggers plasmodesmal closure and defensive signalling, hindering fungal infection progression. **B.** In the absence of the ChEC108-HIPP6 complex, such defence responses are not activated, allowing *C. higginsianum* infection to proceed faster.

### Supplementary Tables

**Table S1. X-ray data collection statistics for ChEC108-HMA1 and ChEC108-HMA2 complex crystals.** Statistics were calculated by *AIMLESS*, implemented in ccp4i2.

| Statistic | | ChEC108-HMA1 | ChEC108-HMA2 |
|---|---|---|---|
| Wavelength (Å) | | 0.6199 | 0.9537 |
| Space group | | $P\ 2_1\ 2_1\ 2_1$ | $P\ 1$ |
| Cell dimensions *a, b, c* (Å) | | 34.91, 76.64, 99.21 | 37.70 60.11, 77.12 |
| Resolution (Å) | Overall | 35.74 - 2.75 | 72.61 – 2.20 |
| | Outer shell | 2.90 – 2.75 | 2.27 – 2.20 |
| $R_{merge}$ (%) | Overall | 18.9 | 11.4 |
| | Outer shell | 91.7 | 85.2 |
| I / $\sigma$I | Overall | 7.9 | 9.0 |
| | Outer shell | 2.1 | 1.3 |
| Completeness (%) | Overall | 100 | 99.4 |
| | Outer shell | 100 | 96.7 |
| Unique reflections | Overall | 7405 | 32007 |
| | Outer shell | 1060 | 2659 |
| Redundancy | Overall | 8.7 | 6.1 |
| | Outer shell | 9.1 | 4.4 |
| CC(1/2) (%) | Overall | 99.3 | 99.6 |
| | Outer shell | 85.6 | 74.8 |

**Table S2. Refinement and model statistics for ChEC108-HMA1 and ChEC108-HMA2 complex crystal structures.** Values were calculated by *REFMAC*, implemented in ccp4i2, or by MolProbity (*). RMSD = root mean square deviation.

| **Statistic** | | **ChEC108-HMA1** | **ChEC108-HMA2** |
|---|---|---|---|
| Resolution range (Å) | | 35.77 - 2.75 | 72.60– 2.20 |
| Number of reflections (all / free) | | 7368 / 379 | 31998 / 1663 |
| $R_{work}$ / $R_{free}$ | | 0.205 / 0.295 | 0.172 / 0.221 |
| Atoms | Protein | 2010 | 4069 |
| | Ion | 1 | 4 |
| | Ligands | 0 | 32 |
| | Water | 19 | 292 |
| B-factors | Protein | 65.94 | 45.44 |
| | Ion | 36.67 | 56.02 |
| | Ligands | - | 52.21 |
| | Water | 38.99 | 44.55 |
| RMSD | Bond lengths (Å) | 0.0045 | 0.0094 |
| | Bond angles (°) | 1.339 | 1.912 |
| Ramachandran * | Favoured | 220 (92.05%) | 477 (96.75%) |
| | Allowed | 18 (7.53%) | 16 (3.25%) |
| | Outliers | 1 (0.42%) | 0 (0%) |
| Clash score * | | 6.47 | 0.85 |
| MolProbity score * | | 2.40 | 1.23 |

**Table S3. $K_d$ values for binding between ChEC108 and HMA domains from different HIPP family members.** $K_d$ values are calculated from ITC experiments displayed in Fig. 3 and Supplementary Fig. S10 and S12.

| HMA domain | Affinity with ChEC108 (nM) | Interaction with $ChEC108^{D164R}$ |
|---|---|---|
| $HIPP6^{HMA1}$ | 1.11 | No |
| $HIPP6^{HMA2}$ | 22.4 | No |
| $HIPP5^{HMA1}$ | 49.3 | No |
| $HIPP34^{HMA}$ | 131 | No |
| $HIPP45^{HMA}$ | 698 | No |
| $HIPP3^{HMA2}$ | 2060 | Not tested |
| $HIPP5^{HMA2}$ | No binding | Not tested |
| $HIPP27^{HMA}$ | No binding | Not tested |

**Table S4. Genotyping primers.** Sequences of primers used to genotype *C. higginsianum* strains (entries 1-10) and *hipp6-3* Arabidopsis (entries 11-12).

| | Primer | Sequence (5'–3') |
|---|---|---|
| 1 | 045_ChEC108_5'Flank_Fw | TTCGCTCAACCCCTAGTACC |
| 2 | 046_ChEC108_3'Flank_Rev | GACCCAGACATCCAGAACCT |
| 3 | 047_trpc_ter_Fw | GTGAATGCTCCGTAACACCC |
| 4 | 048_cos1_pro_Rev | TCGATCTGTTCCTGCCAG |
| 5 | 049_ChEC108_Fw | TGCGACGTATGTTTCCTTCC |
| 6 | 050_ChEC108_Rev | GTCGGCCACTACTACTGGATC |
| 7 | 099_Hyg_Rev | ATGTTGGCGACCTCGTATTG |
| 8 | 100_Hyg_Fw | CTGATCGAAAAGTTCGACAGC |
| 9 | 207_AyB269_Bar_Fw | TGAAGACGAAATGAGCCCAGAACGACGC |
| 10 | 208_AyB270_Bar_Rev | TGAAGACGAAAGCCTAAATCTCGGTGACGGG |
| 11 | 085_CRISPR_HIPP6_Fw | GCACCAGTCCCGGTTAAAAAC |
| 12 | 086_CRISPR_HIPP6_Rev | ACACCAATTCGAACCAAAGACC |

### Legends for Supplementary Datasets

**Dataset S1. GFP- and tdTomato-labelled nuclear counts displayed in Fig. S2.** $35S_{pro}$*:NLS-tdTomato* was transiently expressed in single *N. benthamiana* leaf epidermal cells via low-OD agroinfiltration alongside $35S_{pro}$*:ChEC108-2×GFP* or $35S_{pro}$*:2×GFP* as a control. Mobility of GFP and tdTomato signal was quantified 3 days post-infiltration by counting the number of fluorescent nuclei at transformation sites.

**Dataset S2. Macroscopic lesion measurements displayed in Fig. S4.** 2D area of necrotic lesions induced by WT or *ΔChec108 C. higginsianum* on Col-0 Arabidopsis leaves. Lesion area was normalised by dividing all measurements by the mean area of WT lesions for a given replicate. Dpi = days post inoculation.

**Dataset S3. Appressoria and hyphal counts displayed in Fig. 1 and Fig. S4.** Arabidopsis leaves were infected with WT or *ΔChec108 C. higginsianum* and stained with trypan blue. Infection structures within epidermal cells (appressoria, biotrophic hyphae and necrotrophic hyphae) were visualised using brightfield microscopy and manually quantified. Dpi = days post inoculation.

**Dataset S4. Appressoria and hyphal counts displayed in Fig. S5.** Arabidopsis leaves were infected with WT, *ΔChec108, ΔChec108.ChEC108 or ΔChec108.*$ChEC108^{D164R}$ *C. higginsianum* and stained with trypan blue. Infection structures within epidermal cells (appressoria, biotrophic hyphae and necrotrophic hyphae) were visualised using brightfield microscopy and manually quantified. Dpi = days post inoculation.

**Dataset S5. *N. benthamiana peptides* co-enriched with ChEC108-eGFP.** Peptides identified by mass spectrometry following immunoprecipitation of ChEC108-eGFP or eGFP from *N. benthamiana* leaves. Significance Analysis of INTeractome (SAINT) scores indicate confidence in the assignment of a candidate as an interactor ChEC108. BFDR = bayesian false discovery rate.

**Dataset S6. Macroscopic lesion measurements displayed in Fig 2.** Area of necrotic lesions induced by WT or *ΔChec108 C. higginsianum* on Col-0 or *hipp6-3* Arabidopsis leaves. Lesion area was normalised by dividing all measurements by the mean area of Col-0 lesions for a given replicate. Dpi = days post inoculation.

**Dataset S7. ChEC108-eGFP-labelled nuclear counts displayed in Fig. 4.** $35S_{pro}$*:ChEC108-eGFP* or $35S_{pro}$*:*$ChEC108^{D164R}$*-eGFP* were transiently expressed via bombardment in single Arabidopsis leaf epidermal cells of genotype Col-0 or *hipp6-3*. GFP-labelled nuclei were manually counted at transformation sites.

**Dataset S8. ChEC108-eGFP-labelled cell counts displayed in Fig. 4.** *35S$_{pro}$:ChEC108-eGFP* was transiently expressed in single *N. benthamiana* epidermal cells using low-OD agroinfiltration of leaves transiently expressing *35S$_{pro}$:mCherry-HIPP6* using high-OD agroinfiltration, or *35S$_{pro}$:mCherry* as a control. Mobility of ChEC108-eGFP was quantified by counting the number of fluorescent cells at transformation sites.

**Dataset S9. mScarlet-labelled nuclear counts displayed in Fig. S14.** mScarlet was transiently introduced via bombardment of Col-0, *AtUBQ10$_{pro}$:ChEC108-eGFP* or *AtUBQ10$_{pro}$:ChEC108$^{D164R}$-eGFP* leaves. Mobility of mScarlet was quantified by counting the number of fluorescent nuclei at transformation sites.

**Dataset S10. Differential gene expression analysis to compare Col-0, *ChEC108-eGFP-18* and *ChEC108$^{D164R}$-eGFP-16* transcriptomes.** Comparison of gene expression levels in indicated Arabidopsis transgenic lines or Col-0 controls. baseMean indicates the normalised expression level for a given gene across all samples. Wald statistical tests were conducted using *DESeq2*, and p values adjusted using the Benjamini and Hochberg method. Differentially expressed genes (defined as showing $\log_2$ fold change $\geq |1|$ and Wald test adjusted p value $\leq 0.05$) are highlighted in green (upregulated) or red (downregulated).

**Dataset S11. GO term enrichment analysis of genes upregulated in the presence of ChEC108.** GO terms assigned to differentially expressed genes in lists marked by asterisks in *SI Appendix*, Fig. S15A. Gene ratio = number of genes assigned to each category (k) relative to the number of genes in the input list with at least one biological process (BP) or molecular function (MF) term assigned (n). Background ratio = number of genes in the reference genome annotated with the term (M) relative to the total number of genes in the reference genome with at least one BP or MF term assigned (N). Z scores are descriptive statistics to indicate how far observed counts deviate from expectation. P values were determined by one-sided Fisher's exact tests and adjusted using the Benjamini-Hochberg method.

**Dataset S12. Details of statistical tests.** Outputs of statistical tests conducted to analyse data presented in indicated figure panels. Sample sizes and composition are given in corresponding figure legends. *** indicates $p < 0.001$, ** indicates $p < 0.01$, * indicates $p < 0.05$, ns indicates no significant difference, SE = standard error, df = degrees of freedom, CI = 95% confidence interval.

### SI References


1. C. Engler, R. Kandzia, S. Marillonnet, A one pot, one step, precision cloning method with high throughput capability. *PLoS One* **3** (2008).
2. A. R. Bentham, *et al.*, pOPIN-GG: A resource for modular assembly in protein expression vectors. *bioRxiv* 2021.08.10.455798 (2021).
3. R. J. Lindsay, M. J. Kershaw, B. J. Pawlowska, N. J. Talbot, I. Gudelj, Harbouring public good mutants within a pathogen population can increase both fitness and virulence. *Elife* **5**, e18678 (2016).
4. X. Yan, *et al.*, The transcriptional landscape of plant infection by the rice blast fungus *Magnaporthe oryzae* reveals distinct families of temporally co-regulated and structurally conserved effectors. *Plant Cell* koad036 (2023).
5. R. O'Connell, *et al.*, A Novel *Arabidopsis-Colletotrichum* Pathosystem for the Molecular Dissection of Plant-Fungal Interactions. *Molecular Plant-Microbe Interactions* **17**, 272–282 (2004).
6. K. Labun, *et al.*, CHOPCHOP v3: expanding the CRISPR web toolbox beyond genome editing. *Nucleic Acids Res.* **47**, W171–W174 (2019).
7. B. Castel, L. Tomlinson, F. Locci, Y. Yang, J. D. G. Jones, Optimization of T-DNA architecture for Cas9-mediated mutagenesis in Arabidopsis. *PLoS One* **14**, e0204778 (2019).
8. T. L. Shimada, T. Shimada, I. Hara-Nishimura, A rapid and non-destructive screenable marker, FAST, for identifying transformed seeds of *Arabidopsis thaliana*. *Plant Journal* **61**, 519–528 (2010).
9. E. E. Tee, S. Samwald, C. Faulkner, "Quantification of Cell-to-Cell Connectivity Using Particle Bombardment" in *Plasmodesmata: Methods and Protocols*, Y. Benitez-Alfonso, M. Heinlein, Eds. (Springer US, 2022), pp. 263–272.
10. H. Leung, E. S. Borromeo, M. A. Bernardo, J. L. Notteghem, Genetic analysis of virulence in the rice blast fungus *Magnaporthe grisea*. *Phytopathology* **78**, 1227–1233 (1988).
11. N. J. Talbot, D. J. Ebbole, J. E. Hamer, Identification and characterization of *MPG1*, a gene involved in pathogenicity from the rice blast fungus *Magnaporthe grisea*. *Plant Cell* **5**, 1575–1590 (1993).
12. H. Koga, K. Dohi, O. Nakayachi, M. Mori, A novel inoculation method of *Magnaporthe grisea* for cytological observation of the infection process using intact leaf sheaths of rice plants. *Physiol. Mol. Plant Pathol.* **64**, 67–72 (2004).
13. J. Schindelin, *et al.*, Fiji: an open-source platform for biological-image analysis. *Nat. Methods* **9**, 676–682 (2012).
14. M. Korn, *et al.*, A Genetic Screen for Pathogenicity Genes in the Hemibiotrophic Fungus *Colletotrichum higginsianum* Identifies the Plasma Membrane Proton Pump Pma2 Required for Host Penetration. *PLoS One* **10**, e0125960 (2015).
15. A. Shevchenko, H. Tomas, J. Havli, J. V Olsen, M. Mann, In-gel digestion for mass spectrometric characterization of proteins and proteomes. *Nat. Protoc.* **1**, 2856–2860 (2006).

16. S. Tyanova, T. Temu, J. Cox, The MaxQuant computational platform for mass spectrometry-based shotgun proteomics. *Nat. Protoc.* **11**, 2301–2319 (2016).

17. G. Teo, *et al.*, SAINTq: Scoring protein-protein interactions in affinity purification – mass spectrometry experiments with fragment or peptide intensity data. *Proteomics* **16**, 2238–2245 (2016).

18. J. Lobstein, *et al.*, SHuffle, a novel *Escherichia coli* protein expression strain capable of correctly folding disulfide bonded proteins in its cytoplasm. *Microb. Cell Fact.* **11**, 753 (2012).

19. D. S. Yu, *et al.*, Engineering plant tandem kinase immune receptors expands effector recognition profiles. *Sci. Adv.* **12**, eaef2426 (2026).

20. G. Winter, *xia2*: an expert system for macromolecular crystallography data reduction. *J. Appl. Crystallogr.* **43**, 186–190 (2010).

21. J. Agirre, *et al.*, The *CCP4* suite: integrative software for macromolecular crystallography. *Acta Crystallographica Section D* **79**, 449–461 (2023).

22. P. R. Evans, G. N. Murshudov, How good are my data and what is the resolution? *Acta Crystallographica Section D* **69**, 1204–1214 (2013).

23. A. J. McCoy, *et al.*, *Phaser* crystallographic software. *J. Appl. Crystallogr.* **40**, 658–674 (2007).

24. J. Jumper, *et al.*, Highly accurate protein structure prediction with AlphaFold. *Nature* **596**, 583–589 (2021).

25. M. Mirdita, *et al.*, ColabFold: making protein folding accessible to all. *Nat. Methods* **19**, 679–682 (2022).

26. K. Cowtan, The *Buccaneer* software for automated model building. 1. Tracing protein chains. *Acta Crystallographica Section D* **62**, 1002–1011 (2006).

27. P. Emsley, B. Lohkamp, W. G. Scott, K. Cowtan, Features and development of Coot. *Acta Crystallogr. D Biol. Crystallogr.* **66**, 486 (2010).

28. G. N. Murshudov, *et al.*, *REFMAC5* for the refinement of macromolecular crystal structures. *Acta Crystallographica Section D* **67**, 355–367 (2011).

29. Á. Piñeiro, *et al.*, AFFINImeter: A software to analyze molecular recognition processes from experimental data. *Anal. Biochem.* **577**, 117–134 (2019).

30. F. Krueger, F. James, P. Ewels, E. Afyounian, B. Schuster-Boeckler, FelixKrueger/TrimGalore: v0.6.7-doi via Zenodo. *10.5281/zenodo.5127898* (2018).

31. M. Martin, Cutadapt removes adapter sequences from high-throughput sequencing reads. *EMBnet. J.* **17**, 10–12 (2011).

32. S. Andrews, FastQC: A quality control tool for high throughput sequence data. Available at: https://www.bioinformatics.babraham.ac.uk/projects/fastqc/ (2018).

33. D. Kim, J. M. Paggi, C. Park, C. Bennett, S. L. Salzberg, Graph-based genome alignment and genotyping with HISAT2 and HISAT-genotype. *Nat. Biotechnol.* **37**, 907–915 (2019).

34. P. Ewels, M. Magnusson, S. Lundin, M. Käller, MultiQC: summarize analysis results for multiple tools and samples in a single report. *Bioinformatics* **32**, 3047–3048 (2016).

35. M. I. Love, W. Huber, S. Anders, Moderated estimation of fold change and dispersion for RNA-seq data with DESeq2. *Genome Biol.* **15**, 550 (2014).

36. G. Yu, L.-G. Wang, Y. Han, Q.-Y. He, clusterProfiler: an R Package for Comparing Biological Themes Among Gene Clusters. *OMICS* **16**, 284–287 (2012).

37. M. G. Johnston, C. Faulkner, A bootstrap approach is a superior statistical method for the comparison of non-normal data with differing variances. *New Phytologist* **230**, 23–26 (2021).

38. R. V Lenth, J. Piaskowski, emmeans: Estimated Marginal Means, aka Least-Squares Means. Available at: https://CRAN.R-project.org/package=emmeans (2025).

39. D. Bates, M. Mächler, B. Bolker, S. Walker, Fitting Linear Mixed-Effects Models Using lme4. *J. Stat. Softw.* **67**, 1–48 (2015).

40. A. Kuznetsova, P. B. Brockhoff, R. H. B. Christensen, lmerTest Package: Tests in Linear Mixed Effects Models. *J. Stat. Softw.* **82**, 1–26 (2017).

41. H. Wickham, ggplot2. *WIREs Computational Statistics* **3**, 180–185 (2011).

42. C.-H. Gao, A. Dusa, ggVennDiagram: A "ggplot2" Implement of Venn Diagram. Available at: https://github.com/gaospecial/ggVennDiagram (2026).

43. E. F. Pettersen, *et al.*, UCSF ChimeraX: Structure visualization for researchers, educators, and developers. *Protein Science* **30**, 70–82 (2021).
